# Cell cycle dependent regulation of mRNA export in response to replication stress

**DOI:** 10.64898/2026.09.06.749765

**Authors:** Tobias D. Williams, Kirstyn T. Carey, William B. Hamilton, Linh H. Ngo, Sylvie van Twest, Vincent Murphy, Jieqiong Lou, Andrew B. Das, Emily Deeb, Ching-Seng Ang, Iva Nikolic, Karla J. Cowley, Kaylene J. Simpson, Kristin K. Brown, Elizabeth Hinde, Sean J. Humphrey, Andrew J. Deans, Vihandha O. Wickramasinghe

## Abstract

Nuclear export of mRNA is extensively coupled to transcription and processing of mRNA. How mRNA export is mechanistically regulated remains poorly understood. Here, we uncover a physical checkpoint at nuclear speckles that regulates mRNA export during DNA replication in S-phase of the cell cycle. Unbiased screening approaches identify WEE1 and CHK1 as novel mRNA export regulators. mRNA export complexes are recruited to sites of replication stress in S-phase. WEE1 inhibition prematurely activates CDK1 and PLK1, leading to accumulation of R-loop associated mRNA in large nuclear speckles, with late markers of replication stress present around their periphery. This recruitment is dependent on CDK1 activity. Phosphorylation of ALYREF by CDK1 and PLK1 regulates nuclear speckle accumulation of mRNA following replication stress. mRNA export factors including ALYREF are subsequently mis-localised from these speckles, preventing nuclear export of R-loop associated mRNA. Thus, WEE1, CDK1 and PLK1 enforce a cell cycle regulated checkpoint at nuclear speckles that serves to protect the cell from major sources of genome instability by ensuring that R-loop associated mature mRNA is not exported to the cytoplasm.

---

Nuclear export of messenger RNA (mRNA) is a critical step in the gene expression pathway and is coupled to transcription, splicing and polyadenylation of mRNA^1^. In humans, this requires export adaptors such as the TREX (TRanscription-EXport)^2–4^ and TREX-2^5–8^ complexes, which recognise mRNAs at sites of transcription and processing^9–18^. One cellular compartment that is intimately linked with mRNA processing and export is the nuclear speckle. Nuclear speckles are composed of RNA splicing and export factors as well as mature poly(A)+RNA^19–22^. Indeed, in human nuclei, the majority of poly(A)+RNA co-localises with nuclear speckles, however the functional role of poly(A)+RNA in speckles remains divisive^23,24^. Some mRNAs may transit through nuclear speckles prior to export^25,26^ and speckles have been proposed to be involved in quality control of mRNAs^27,28^ and more recently, in regulation of mRNA export during transcriptional stress^11^. While nuclear speckles do not contain DNA, recent observations suggest that a number of highly expressed active genes are recruited to the periphery of nuclear speckles for transcription and processing ^9,24,29–33^. Whether the nuclear speckle plays other cellular functional roles remains unknown.

Transcription, processing and nuclear export of mRNA occurs throughout the cell cycle. However, during S-phase, replication of DNA occurs on the same template as transcription by the RNA polymerase II machinery, and there is the potential for conflicts between the DNA replication and transcription machineries^34^. One potent barrier to DNA replication fork progression are R-loops, a transcription intermediate containing DNA hybridised to the transcribed RNA strand (DNA:RNA hybrid), along with displaced ssDNA (reviewed in ^35,36–39^). Interestingly, R-loops have recently been detected in the cytoplasm and can elicit an immune response^40–42^, raising the possibility that there may be an active mechanism that prevents their export from the nucleus into the cytoplasm.

Here, through unbiased screening and phosphoproteomic approaches, we uncover a physical checkpoint mediated by WEE1, CDK1 and PLK1 that regulates mRNA export in response to replication stress during S-phase. Our findings demonstrate that mRNA export is mechanistically regulated during the cell cycle and highlight a previously unidentified and important functional role for nuclear speckles in processing of unresolved R-loops that contain poly(A)+RNA. We propose that this checkpoint serves to protect the cell from major sources of genome instability by ensuring that R-loop associated mature mRNA is not exported to the cytoplasm.

## Results

### CRISPR and siRNA screens identify cell cycle DNA damage kinases WEE1 and CHK1 as mRNA export regulators

While the protein components of the mRNA export machinery have been well characterised, how mRNA export is mechanistically regulated remains poorly understood. For example, is mRNA export regulated by phosphorylation and if so, which kinases function in this process? To address this question, we used two independent technologies, namely siRNA and CRISPR-Cas9 to perform a semi-automated high content screen against the human kinome (Figure 1A-C). We then assessed whether depletion or deletion of human kinases resulted in an increase in the nuclear/cytoplasmic (N/C) ratio of poly(A)+RNA. RNA export defects were quantified by measuring the N/C ratio of poly(A)+RNA across thousands of individual cells, whereby an increase indicates an mRNA export block (Figure S1A, B). As positive controls, we included known components of the TREX (ALYREF), TREX-2 (GANP) complexes and NXF1, whose depletion or deletion resulted in an inhibition of mRNA export, as expected (Figure 1B,C and Figure S1A,B). Unexpectedly, WEE1 and CHK1, which both function to regulate CDK1 activity in the S-phase of the cell cycle and regulate the G_2_/M DNA damage checkpoint, were identified as potential regulators of mRNA export in both CRISPR and siRNA screens (Figure 1B,C). We next validated these hits using three independent methods, namely CRISPR-Cas9 deletion, depletion by independent siRNAs and chemical inhibition of WEE1 and CHK1, all of which inhibited the nuclear export of poly(A)+RNA (Figures 1D-J and S1A,B). Strikingly, during our screening and validation experiments, we consistently observed that depleting or inhibiting CHK1 or WEE1 resulted in a subset of cells exhibiting an mRNA export defect, with WEE1 depletion/inhibition generally resulting in a higher percentage of cells displaying a defect (Figures 1D-H and S1A). Indeed, when we temporally assessed WEE1 inhibition, the percentage of cells displaying an increase in the nuclear/cytoplasmic ratio of poly(A)+RNA increased with time of inhibition (Figure 1I-J). Interestingly, these cells displayed massively enlarged nuclear poly(A)+RNA foci when compared to both controls and depletion of traditional mRNA export factors (Figure 1D,E). The cells with enlarged nuclear mRNA foci were the driver of the increase in nuclear/cytoplasmic ratio with the majority of cells having normal distributions of poly(A)+RNA (Figure 1G). We confirmed that these enlarged foci were indeed nuclear speckles, using poly(A)+RNA fluorescent in situ hybridisation (FISH) combined with immunofluorescence of nuclear speckle marker SC35 (Figure S1C). Importantly, this differs from the universal defect exhibited when traditional mRNA export factors are perturbed (Figures 1D,E and S1A), raising the possibility that cells with an mRNA export defect following WEE1 inhibition are in a particular phase of the cell cycle.

**Figure 1.**
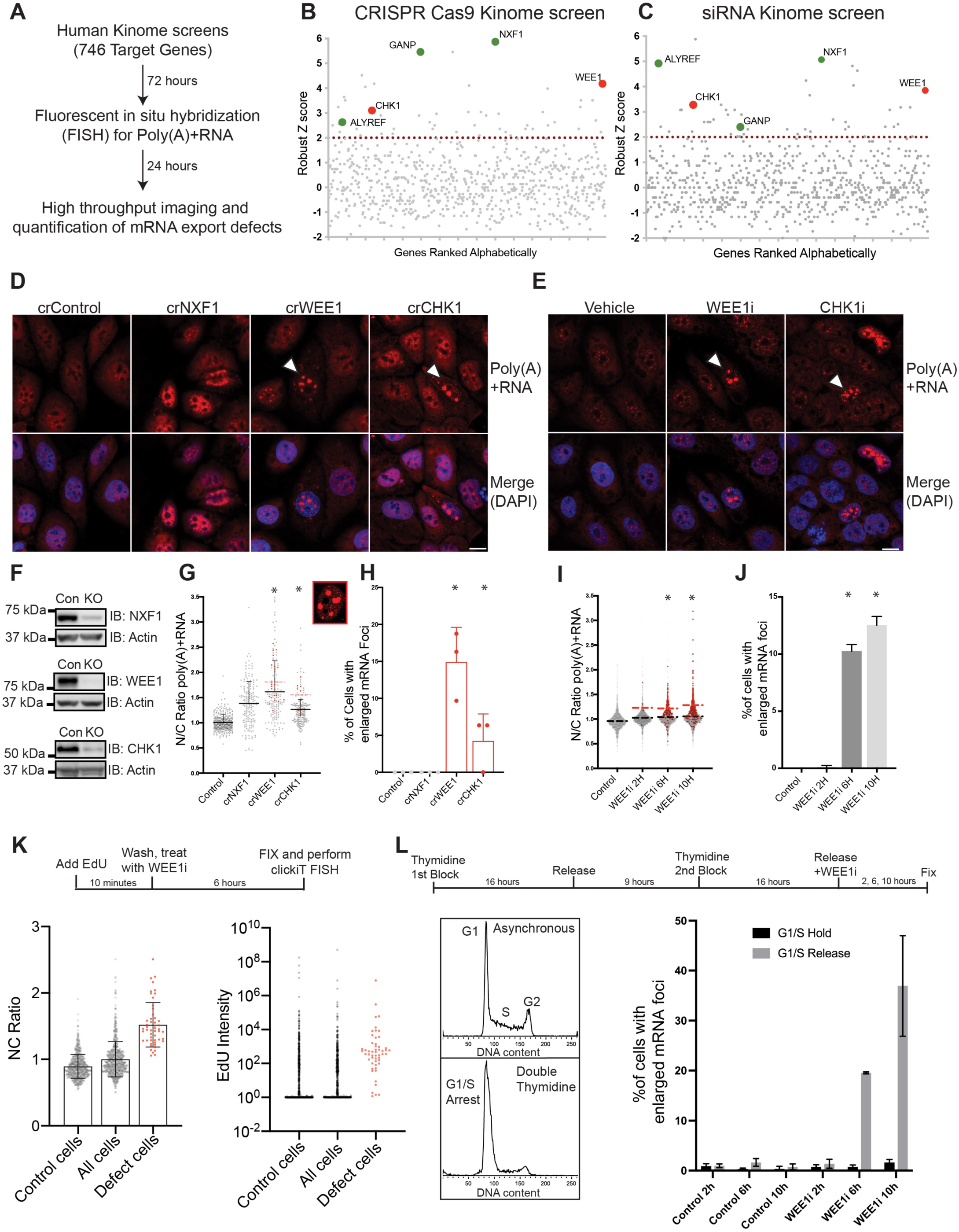
CRISPR and siRNA screens both identify cell cycle and DNA damage regulating kinases WEE1 and CHK1 as mRNA export regulators. (**A)** Workflow for high throughput CRISPR and siRNA kinome screens to identify novel mRNA export regulators. Nuclear/Cytoplasmic (N/C) ratio of poly(A)+RNA was assessed following individual depletion or deletion of 675 protein kinases. (**B,C)** Z-scores for individual genes are shown for the CRISPR-Cas9 screen (**B**) and siRNA screen (**C**) performed in (**A)**. Positive controls are enlarged and highlighted in green and novel regulators CHK1 and WEE1 identified by the screen are enlarged and highlighted in red. Z score of 2 is indicated by the dotted line with those genes above the line considered as significant. (**D)** Representative confocal images of RNA-FISH for poly(A)+RNA following perturbation of control, NXF1, CHK1 and WEE1 via CRISPR-Cas9 mediated deletion. White arrows indicate a subpopulation of cells with enlarged poly(A)+ RNA foci. (**E)** Representative confocal images of RNA-FISH for poly(A)+RNA following chemical inhibition of CHK1 or WEE1. White arrows indicate a subpopulation of cells with enlarged poly(A)+RNA foci. (**F)** Knockout efficiency of NXF1, WEE1 and CHK1 was monitored by western blotting with indicated antibodies. (**G)** Analysis of N/C ratio of poly(A)+RNA of (**D**) with a minimum of 50 cells analysed per experimental repeat (n=3). Cells with enlarged poly(A)+ RNA foci are highlighted in red. (**H)** Number of cells with enlarged poly(A)+RNA foci identified in (**D**) represented as a percentage. (**I)** Analysis of N/C ratio of poly(A)+RNA of (**E**) with a minimum of 50 cells analysed per experimental repeat (n=3). Cells with enlarged poly(A)+ RNA foci are highlighted in red. (**J)** Number of cells with enlarged poly(A)+RNA foci identified in (**I**) represented as a percentage across 3 individual experimental repeats. (**K**) Cells need to progress through S-phase to acquire the specific mRNA export defect. N/C ratio of poly(A)+RNA and EdU nuclear intensity were assessed following labelling of S-phase cells just prior to treatment with a WEE1 inhibitor using EdU and Click iT-RNAFISH. Cells with nuclear poly(A)+RNA foci one standard deviation larger than the mean of the control are plotted separately and represented in red. (**L**) Cells were synchronised in S-phase using a double thymidine block and either released or held at the G_1_/S boundary and treated with a WEE1 inhibitor for the indicated times. Flow cytometry analysis of DNA content (left panel) and analysis of cells with nuclear poly(A)+RNA foci one standard deviation larger than the mean of the control represented as a percentage (right panel) are indicated. Graphs represent the mean + SD or the mean +/− SD with * indicating p <0.05 (Students t test). Cells with nuclear poly(A)+RNA foci 1 standard deviation larger than the mean of the control are represented in red, cells with normal patterns of poly(A)+RNA in are grey.

### WEE1 signalling and replication stress regulate nuclear export of mRNA in S-phase

To determine whether the cells with enlarged nuclear speckles are linked with cell cycle stage, we used quantitative image-based cytometry combined with poly(A)+RNA FISH to simultaneously analyse the mRNA localisation profiles of G_1_, S and G_2_ cells following WEE1 inhibition (Figure S1D-F). Actively replicating cells were marked at the time of harvest using a synthetic nucleotide derivative, 5-Ethynyl-2’deoxyurdine (EdU). As observed previously, WEE1 inhibition resulted in a progressive decrease over time of EdU incorporation (Figure S1D-F) indicating a slowing of DNA replication, or replication stress ^43^. Interestingly, the cells with enlarged nuclear mRNA foci following WEE1 inhibition exhibited reduced levels of EdU incorporation and DNA intensity, consistent with them undergoing replication stress in S-phase (Figure S1D-F). To determine whether WEE1 inhibited cells need to progress through S-phase in order to acquire the mRNA export defect, we next pulsed cells with EdU just prior to WEE1 inhibition to mark actively replicating cells. These experiments demonstrated that cells with enlarged mRNA foci have increased EdU intensity in contrast to those with no mRNA export defect, indicating that they are in S-phase at the time of treatment (Figure 1K).

These experiments suggest that the reason that only a subset of WEE1 inhibited cells have an mRNA export defect is because they are in different phases of the cell cycle at the time of inhibition. Indeed, cells that are synchronised at the G_1_/S boundary using a double thymidine block and subsequently released into S-phase exhibited a large increase in the percentage of cells with an mRNA export block in response to WEE1 inhibition (Figure 1L). Critically, cells held at the G_1_/S boundary and treated with WEE1 inhibitors had a normal distribution of poly(A)+ RNA, indicating that the cells need to enter S-phase in order to acquire the mRNA export defect (Figure 1L). Taken together, these two approaches, namely imaging based cytometry and cell synchronisation, demonstrate that aberrant WEE1 signalling regulates nuclear export of mRNA in S-phase.

### mRNA export complexes are recruited to sites of replication stress

Our observations thus far raised the intriguing possibility that cells may have an active mechanism that regulates mRNA export during DNA replication in S-phase. Nuclear export of mRNA is extensively coupled to transcription and processing of mRNA throughout the cell cycle, however during S-phase, replication of DNA occurs on the same template as transcription by the RNA polymerase II machinery. This raises the important question of what happens to transcription and mRNA export in S-phase when collisions between the replication and transcription machineries may occur. Indeed, our data suggested that cells with enlarged nuclear mRNA foci following WEE1 inhibition are undergoing replication stress in S-phase (Figure S1D-F).

First, we asked whether mRNA export factors play a direct role in replication stress-coupled mRNA processing. We determined whether TREX components could interact with replication protein A (RPA) following induction of replication stress with camptothecin, a topoisomerase 1 (TOP1) inhibitor, or WEE1 inhibition. RPA is a single-stranded DNA binding protein that co-ordinates the recruitment and exchange of genome maintenance factors during DNA replication stress ^44^. We enriched for RPA bound to chromatin in the nucleus and confirmed that our structure bound fraction was enriched for Histone H3, but not Exportin-5 (XPO5) which was instead enriched in the nuclear soluble fraction (Figure 2A). Endogenous TREX components including THOC2 and ALYREF were co-immunoprecipitated from structure-bound nuclear extracts of CAL51 cells undergoing replication stress by antibodies against RPA as was mRNA transport factor NXF1 (Figure 2B), suggesting that they can interact *in vivo.* The interaction was strongest in cells treated with WEE1 inhibitors or both WEE1 inhibitors and camptothecin, where immunoprecipitated RPA was phosphorylated at serine residues 4 and 8 (S4/S8), a known marker of replication stress (Figure 2B) ^45–47^. PRP19, previously shown to associate with RPA ^48^, was also co-immunoprecipitated from this fraction (Figure 2B). Next, we assessed whether TREX component ALYREF was enriched at sites of replication stress in cells, marked by phosphorylated RPA. We observed that in camptothecin treated cells, ALYREF co-localised with a proportion of phosphorylated RPA throughout the nucleus (Figure 2C). Moreover, in WEE1 inhibited cells and camptothecin and WEE1 inhibitor treated cells with small nuclear speckles, we again observed co-localisation with phosphorylated RPA (Figure 2C). Taken together, these results suggest that TREX components can be recruited to sites of replication stress.

**Figure 2.**
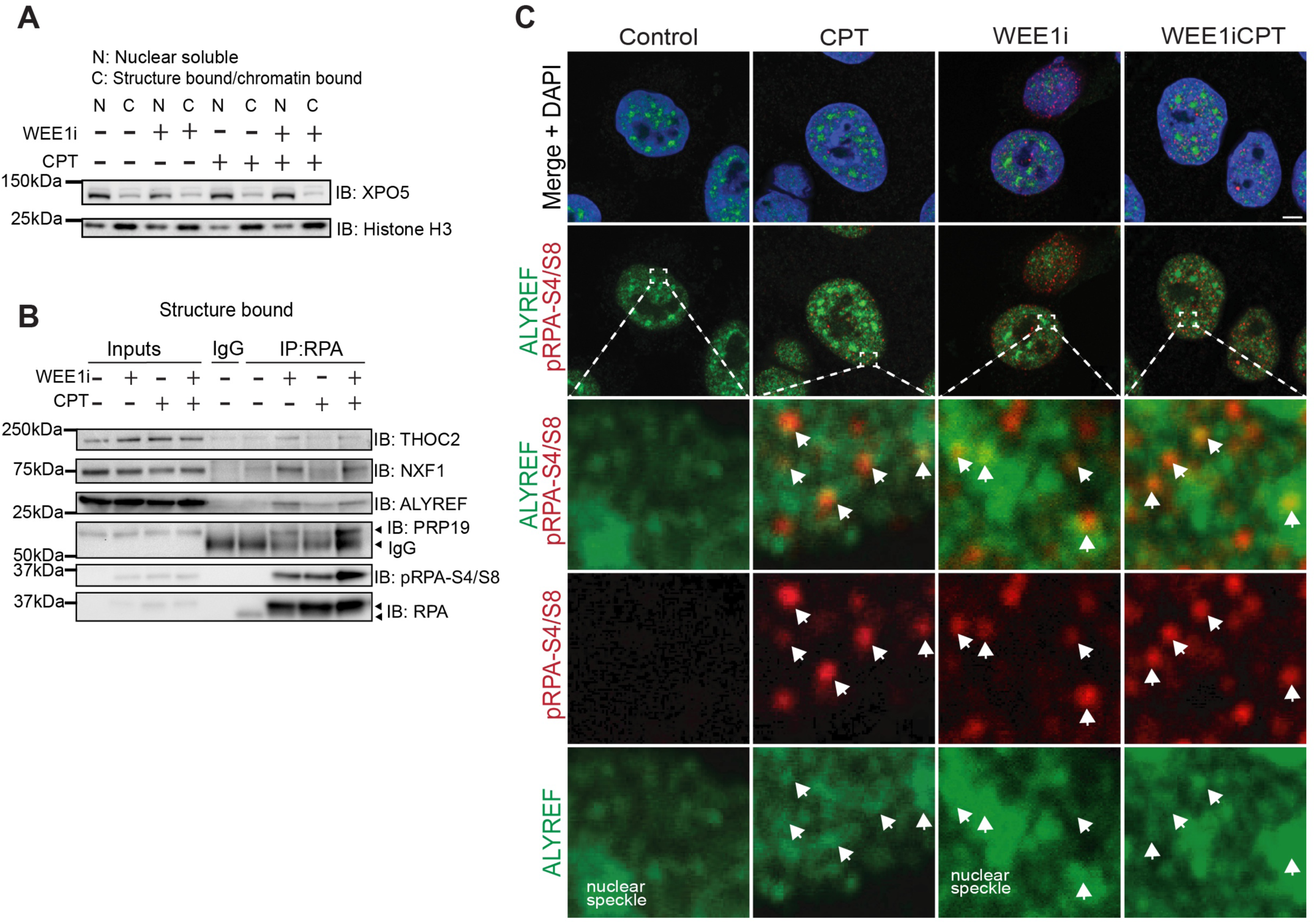
mRNA export factors are recruited to sites of replication stress. **(A)** Efficiency of nuclear soluble and structure bound/chromatin bound fractionation was monitored by western blotting with indicated antibodies. Samples were prepared from S-phase synchronised cells treated with the indicated inhibitors. (**B**) NXF1, THOC2 and ALYREF interact with RPA following replication stress. Endogenous RPA was immunoprecipitated from structure-bound nuclear extract and interacting proteins were analysed by western blotting with the indicated antibodies. PRP19 is shown as a positive control. (**C**) ALYREF co-localises in part with phosphorylated RPA following replication stress. Representative confocal microscopy images of dual immunofluorescence for replication stress marker pRPA-S4/S8 and ALYREF. Cells with similar sized nuclear speckles that best represented n of 3 experiments were chosen to represent an early phenotype. For all experiments, cells were first synchronised in S-phase via double thymidine block, released and treated with the indicated inhibitors for 6 hours. White arrows indicate colocalisation.

To understand the potential contribution of replication stress to prevention of mRNA export in S-phase, we next induced replication stress and observed the localisation and levels of RPA, phosphorylated at serine residues 4 and 8 (S4/S8) and poly(A)+RNA. Interestingly, we saw differential effects on mRNA export in response to different replication stress inducers. Induction of replication stress with camptothecin resulted in an mRNA export defect in smaller foci reminiscent of transcription factories (Figure 3A-C). In contrast, cells treated with the ribonucleotide reductase inhibitor hydroxyurea (HU) had a normal distribution of poly(A)+RNA (Figure S2A-D). Importantly, both inhibitors had similar levels of replication stress as indicated by phosphorylated RPA staining (Figures 2A-B and S2A-D), indicating the mRNA export block was specific to topoisomerase 1 inhibition.

**Figure 3:**
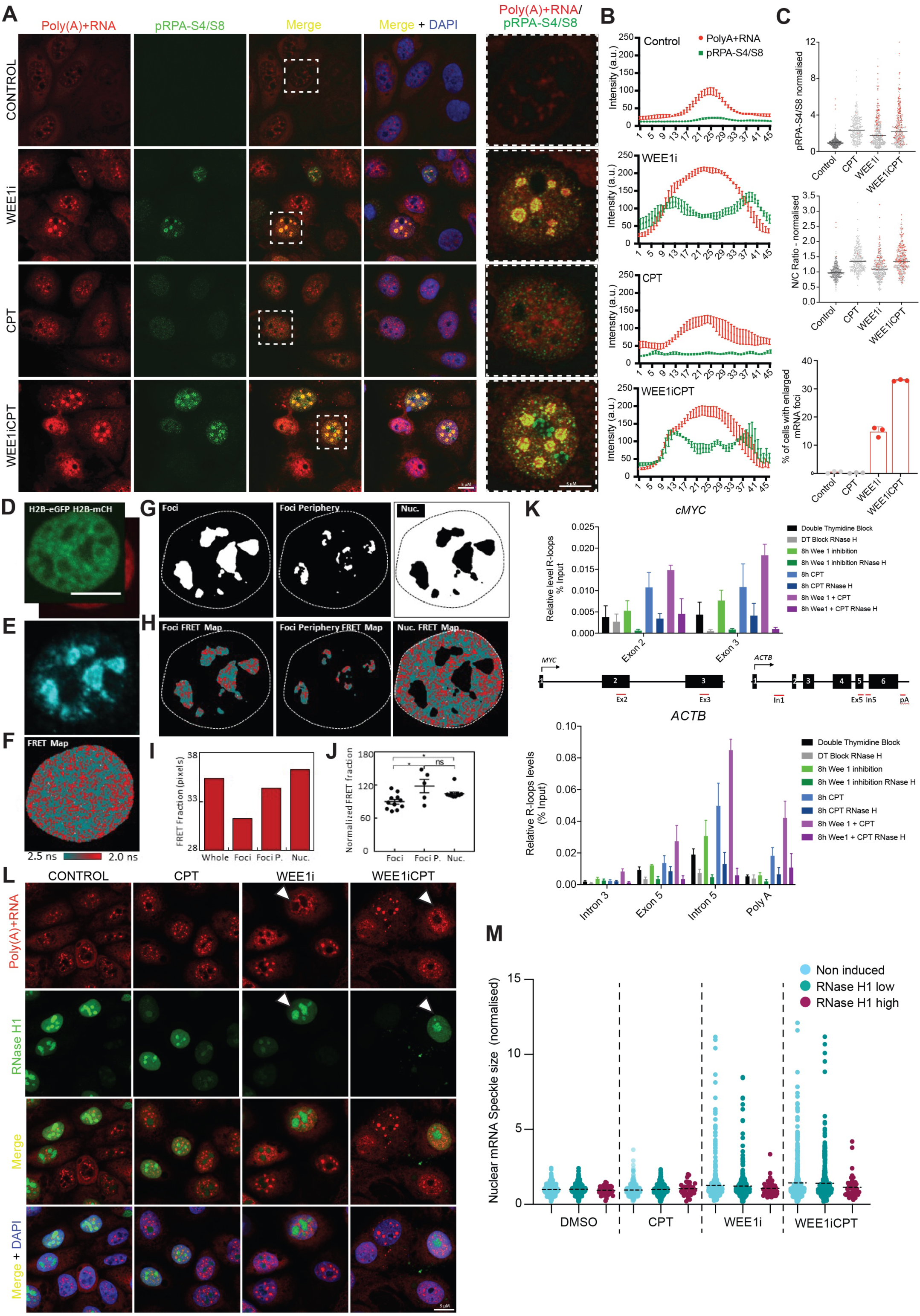
Enlarged mRNA speckles are composed of ssDNA, poly(A)+RNA and R-loops. **(A)** Phosphorylated RPA localises to the periphery of enlarged nuclear speckles following replication stress. Representative confocal microscopy images of FISH for poly(A)+RNA and IF for pRPA-S4/S8 for the indicated treatments are shown. Individual cells from the merge channel are selected and displayed at a higher magnification, those selected are indicated by the white box. **(B**) Line scan analysis of fluorescent intensity of (**A**) performed on individual speckles of the same size. Lines represent the mean intensity across 5 speckles selected from three individual experiments with error bars showing SEM. (**C**) Analysis of nuclear intensity of pRPA-S4/S8, N/C ratio and percentage of cells with enlarged nuclear speckles for the indicated treatments. Cells with nuclear speckles greater than one standard deviation above the control cells are highlighted in red. A minimum of 50 individual cells were analysed per experimental repeat (n=3). **(D-E)**,CAL51 cell nucleus co-expressing the histone FRET pair H2B-eGFP and H2B-mCherry (**D**). SRRM2 immunofluorescence of this nucleus is shown in (**E**). **(F**) FLIM map of the cell presented in (D,E) pseudo-coloured to report histone FRET (red pixels) versus non-FRET (teal pixels). **(G**) Masks based on SRRM2 IF presented in panel (E) that select chromatin inside (left), at edge (middle), and outside (right) of enlarged poly(A)+RNA nuclear speckles. **(H)** Fractions of histone FRET (compact chromatin) in the histone FRET map presented in panel (**G**). **(I**) Quantification of fraction of histone FRET inside, on the edge and outside of late stage enlarged poly(A)+RNA nuclear speckles across multiple cells (n= 11 cells, two biological replicates). (**J**) Scatter plot showing normalized individual cell value from I with standard error of mean (SEM). * *p* < 0.05 (paired *t*-test). **(K)** DRIP qPCR assessment of R-loop formation in cMYC and ACTB genes in the indicated genomic regions. Graphs represent the mean of four independent experiments with error bars showing SEM. (**L,M**) Cells expressing high levels of RNase H1 do not form enlarged nuclear speckles. FISH for polyA(+)RNA and IF for RNase H1 (**L**). Cells were synchronised in S-phase via double thymidine block, released and treated with the indicated inhibitors for 6 hours. Cells with high expression of RNase H1 and small nuclear speckles are indicated with the white arrow heads. Nuclear speckle size was quantified in cells with/without RNAse H1 induction (**M**). Graph represents data from 4 independent experiments with a minimum of 50 cells per experiment. High RNase H1 was defined by those cells with the top 15% of GFP fluorescent intensity.

When we combined WEE1 inhibition with campthothecin or hydroxyurea treatment, we found that replication stress potentiates the mRNA export defects resulting from WEE1 inhibition. Thus, cells treated with camptothecin and WEE1 inhibitors had a large increase in the percentage of cells with enlarged nuclear speckles, which correlated with increased RPA-pS4/S8 intensity (Figure 3A-C), while treating with hydroxyurea prevented their formation (Figure S2A-D). Strikingly, phosphorylated RPA was localised at the periphery of these enlarged nuclear speckles (Figure 3A, right panels and Figure 3B, line scans). As phosphorylated RPA marks damaged single-stranded DNA (ssDNA) ^44^, this result suggests that the enlarged poly(A)+RNA nuclear speckles are encapsulated by ssDNA bound by RPA.

To assess how chromatin architecture is influenced by the enlarged poly(A)+RNA nuclear speckles, we used phasor approach to fluorescence lifetime imaging microscopy (FLIM) of Förster resonance energy transfer (FRET) between fluorescently labelled histones to directly measure the chromatin network architecture at nanometre resolution in intact nuclei (Figure S3A-E). This technique enabled us to detect whether there were any changes in this structural framework following camptothecin treatment and WEE1 inhibition (Figure S3F-H). No nuclear-wide chromatin architecture changes were observed in response to camptothecin treatment and WEE1 inhibition (Figure S3I). To test whether the local chromatin architecture in the enlarged poly(A)+RNA nuclear speckles is altered, we coupled histone FLIM-FRET with immunofluorescence against poly(A)+RNA nuclear speckle marker SC35 (Figure 3D-F). We next generated an immunofluorescence intensity-based mask that enabled histone FRET analysis of chromatin architecture inside, at the periphery of, and outside of enlarged poly(A)+RNA nuclear speckles (Figure 3G,H). Through quantification of the fraction of histone FRET pixels within the FLIM map that occupy inside, at the periphery, and outside of enlarged poly(A)+RNA nuclear speckles (Figure 3I), our results suggested that chromatin architecture at enlarged poly(A)+RNA nuclear speckles is more ‘open’ compared to the rest of the nucleus (Figure 3J). Interestingly, the periphery of the enlarged poly(A)+RNA nuclear speckles was significantly more compact compared to the interior of poly(A)+RNA nuclear speckles (Figure 3J). This may be because we primarily observe phosphorylated RPA that marks damaged single-stranded DNA at the periphery of the enlarged poly(A)+RNA nuclear speckles. The local chromatin structure inside of the normal poly(A)+RNA nuclear speckles and the early stages of enlarged poly(A)+RNA nuclear speckles were also more ‘open’ compared to the rest of the nucleus (Figure S3J-L). Importantly, the more ‘open’ chromatin structure that we observed in the poly(A)+RNA nuclear speckles region is not a low histone concentration artefact, as chromatin architecture at nucleoli, another region with low histone concentration, was less ‘open’ than the enlarged poly(A)+RNA nuclear speckles (Figure S3M-P).

### Enlarged mRNA speckles are composed of ssDNA, poly(A)+RNA and RNA/DNA hybrids

Our results raised the possibility that the increase in nuclear speckle size and changes in chromatin architecture that we observed in response to replication stress and WEE1 inhibition may be driven at least in part by the re-localisation of RPA bound ssDNA and TREX components from sites of replication stress to nuclear speckles. Indeed, TREX components are observed both at RPA foci and in nuclear speckles (Figure 2C). Importantly, these enlarged nuclear speckles contain an abundance of mature polyadenylated mRNA, raising the question of whether these foci actually contain R-loops, a transcription intermediate containing DNA hybridised to the transcribed RNA strand (DNA:RNA hybrid), along with displaced ssDNA reviewed in ^35,36,37^. R-loop formation at highly transcribed regions can be prevented by TOP1 ^49,50^, which relaxes co-transcriptionally generated negative supercoiling of DNA behind elongating RNA polymerases. If unresolved, this may lead to local unwinding of DNA strands, increasing the probability that RNA hybridizes to the DNA. As both RPA and TOP1 are linked with preventing or resolving pathological R-loops, this suggested that the formation of these foci is potentially linked with WEE1 signalling during replication stress induced by replication-transcription conflicts. We therefore determined whether R-loop levels were increased following WEE1 inhibition. First, we performed DRIP-qPCR using the S9.6 antibody, which recognises R-loops. As this antibody is known to also bind to double stranded RNA, we utilised RNase H, which specifically degrades R-loops to confirm the specificity of the signal. We found that R-loop levels were increased following WEE1 inhibition and by the addition of TOP1 inhibitor camptothecin and this was RNaseH dependent (Figure 3K). Next, we assessed whether the increase in nuclear mRNA speckle size that we observed was due to increased levels of R-loops. As the S9.6 antibody has been shown to be problematic for use in immunofluorescence experiments ^51,52^, we generated a cell line with inducible RNase H1 and determined whether RNase H1 induction could alter nuclear speckle size following replication stress and WEE1 inhibition. We found that in WEE1 inhibited or WEE1 and TOP1 inhibited cells expressing high levels of RNase H1, nuclear speckle size was reduced to basal levels (Figure 3L,M). While we did observe a number of cells with enlarged mRNA foci following WEE1 inhibition, these cells had lower levels of RNase H1 (Figure 3L). Taken together, these results suggest that enlarged nuclear speckles that form following severe replication stress contain mature polyadenylated mRNA hybridised to DNA and the displaced ssDNA.

### mRNA export factors such as ALYREF are phosphorylated by CDK1 and PLK1 in response to replication stress

We next sought to confirm whether the kinase responsible for regulation of mRNA export during replication stress was WEE1 or kinase/s downstream from WEE1. WEE1 and CHK1 maintain control of CDK1 activity during normal DNA replication in S-phase ^53^. Thus, WEE1 normally phosphorylates CDK1 at tyrosine residue 15 to inhibit its activity ^54^ and its inhibition or depletion induces premature activation of CDK1 in G_1_ and S phase cells, leading to replication stress ^43,53,55^. As CDK1 is linked with induction of replication stress following WEE1 inhibition, we aimed to quantify temporal changes in protein phosphorylation following WEE1 inhibition, CDK1 inhibition and/or induction of replication stress with TOP1 inhibition. We decided to use an unbiased temporal phosphoproteomics approach ^56^ to identify changes in phosphorylation patterns following replication stress. Cells were harvested 30 mins, 2 and 6 hours post treatment with either camptothecin, CDK1 or WEE1 inhibitors or combinations thereof. We identified 28,764 phosphorylation sites across the experiment with 21,117 identified in all four replicates (Figure 4A). Global phosphorylation analysis indicated a number of cellular effects consistent with WEE1 inhibition, including reduced inhibitory phosphorylation of CDK1 at tyrosine residue 15 within 30 minutes of treatment (Figure S4A), accompanied by increased phosphorylation of CDK1 and CDK2 substrates 2 hours post treatment, which returned to baseline 6 hours post treatment (Figure 4B). WEE1 inhibition has been reported to result in an increased interaction between the G_2_/M kinase PLK1 and CyclinA2-CDK, resulting in premature PLK1 activation ^57^. In agreement, we observed increased activity of PLK1 as evidenced by an increase in phosphorylation of PLK1 substrates, but this activation was temporally separated from CDK activation as the increase was only observed 6 hours after WEE1 inhibition (Figure 4B). Furthermore, CDK1 inhibition led to a global decrease in CDK1 activity and addition of camptothecin led to activation of DNA damage kinases CHK1, ATM and ATR ^58,59^, consistent with its well established role in inducing replication stress (Figure 4B).

**Figure 4:**
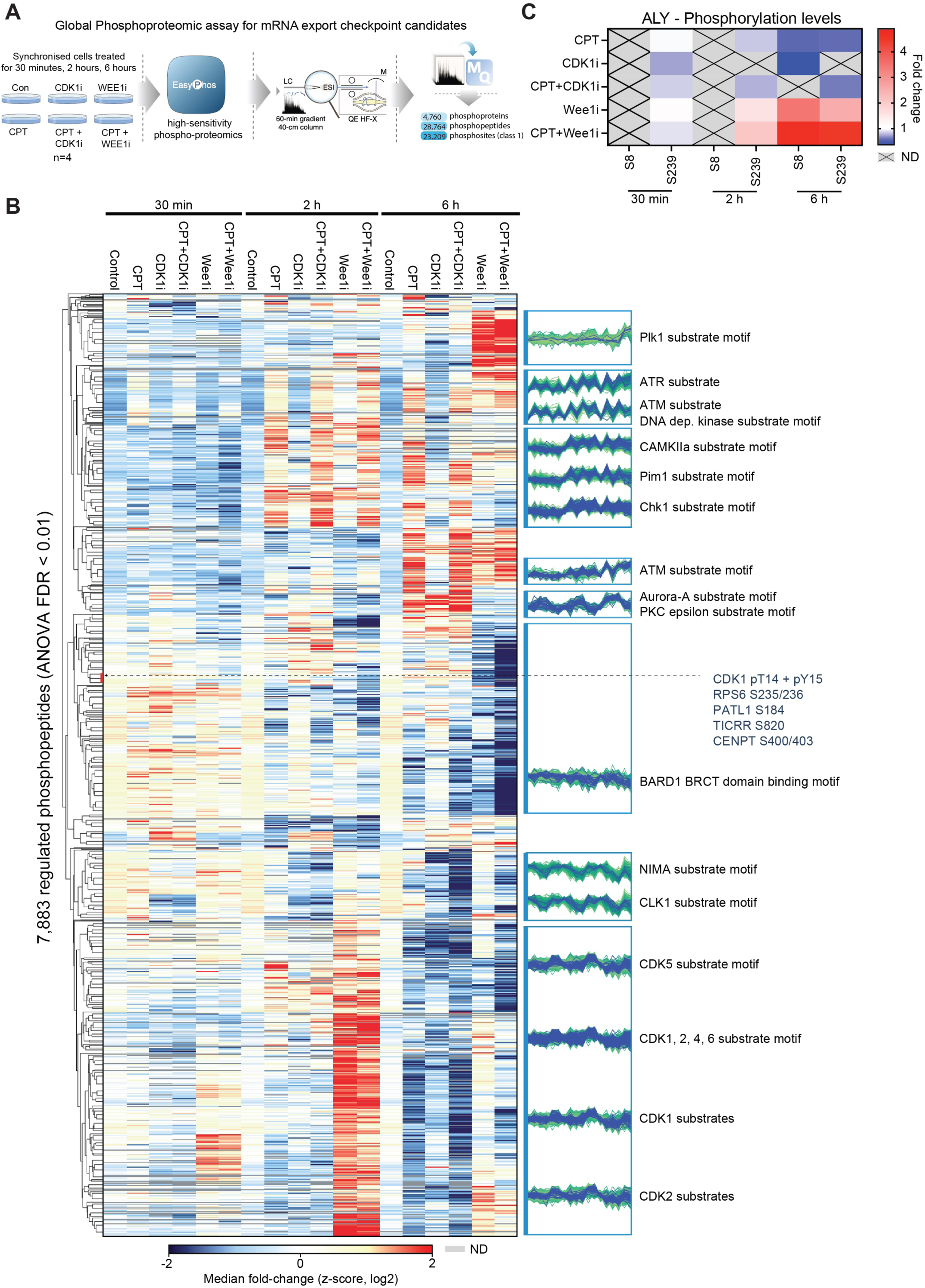
mRNA export factors including ALYREF are phosphorylated in response to replication stress. **(A**) Workflow for high throughput EasyPhos global phosphoproteomic assay to assess phosphoproteomic profiles of cells progressing through S-phase following the indicated treatments. (**B)** Heat map of global phosphorylation profiles of cells under replication stress with or without the indicated kinase inhibition across each timepoint. Colours represent a fold change of phosphorylation-normalised to control (n=4). (**C)** Heat map of phosphorylation changes of mRNA export factor ALYREF identified during the EasyPhos experiment in **(A,B)**. Peptides that were not detected (ND) in particular timepoints are represented in grey.

Further analysis of this large-scale proteomic dataset revealed that several mRNA export factors, including THOC2, THOC5, THOC7, GANP, UAP56 and ALYREF are phosphorylated during replication stress in a manner that is altered by WEE1 inhibition (Figure 4C and Figure S4B). Most interestingly, ALYREF was phosphorylated at distinct residues in its N and C-termini that have been proposed to be critical for its role in mRNA export ^60–62^. Under conditions of replication stress and CDK1 inhibition, when nuclear speckles are small, reduced phosphorylation occurred at these residues. Conversely, under conditions of WEE1 inhibition, when nuclear speckles are large, a significant increase in phosphorylation was observed (Figure 4C). This raised the question as to whether the formation of enlarged nuclear speckles with increased R-loops is driven by cell cycle kinase regulation of ALYREF on serine 8 or serine 239. To identify putative responsible kinases, we compared sequences surrounding ALYREF S8 and S239 to known kinase substrate motifs and identified PLK1, CDK1 and casein kinase 2 (CK2) as candidates. Importantly, PLK1, WEE1 and CDK1 are also functionally linked in the S/G_2_ phase of the cell cycle and temporal patterns of global PLK1/CDK1 activity identified in the phospho-proteomic analysis matched with the timing of ALYREF phosphorylation (Figure 4C). Thus, we next performed *in vitro* CDK, CK2 and PLK1 kinase assays with full length ALYREF protein and analysed phosphorylation by autoradiography and mass spectrometry (Figure 5A,B). These experiments indicated that ALYREF is phosphorylated by PLK1 at serine 8 and by CDK at serine 239 (Figure 5B). These were the only phosphorylation sites observed by mass spectrometry and no phosphorylation was observed when the promiscuous kinase CK2 was used (Figure S5A).

**Figure 5:**
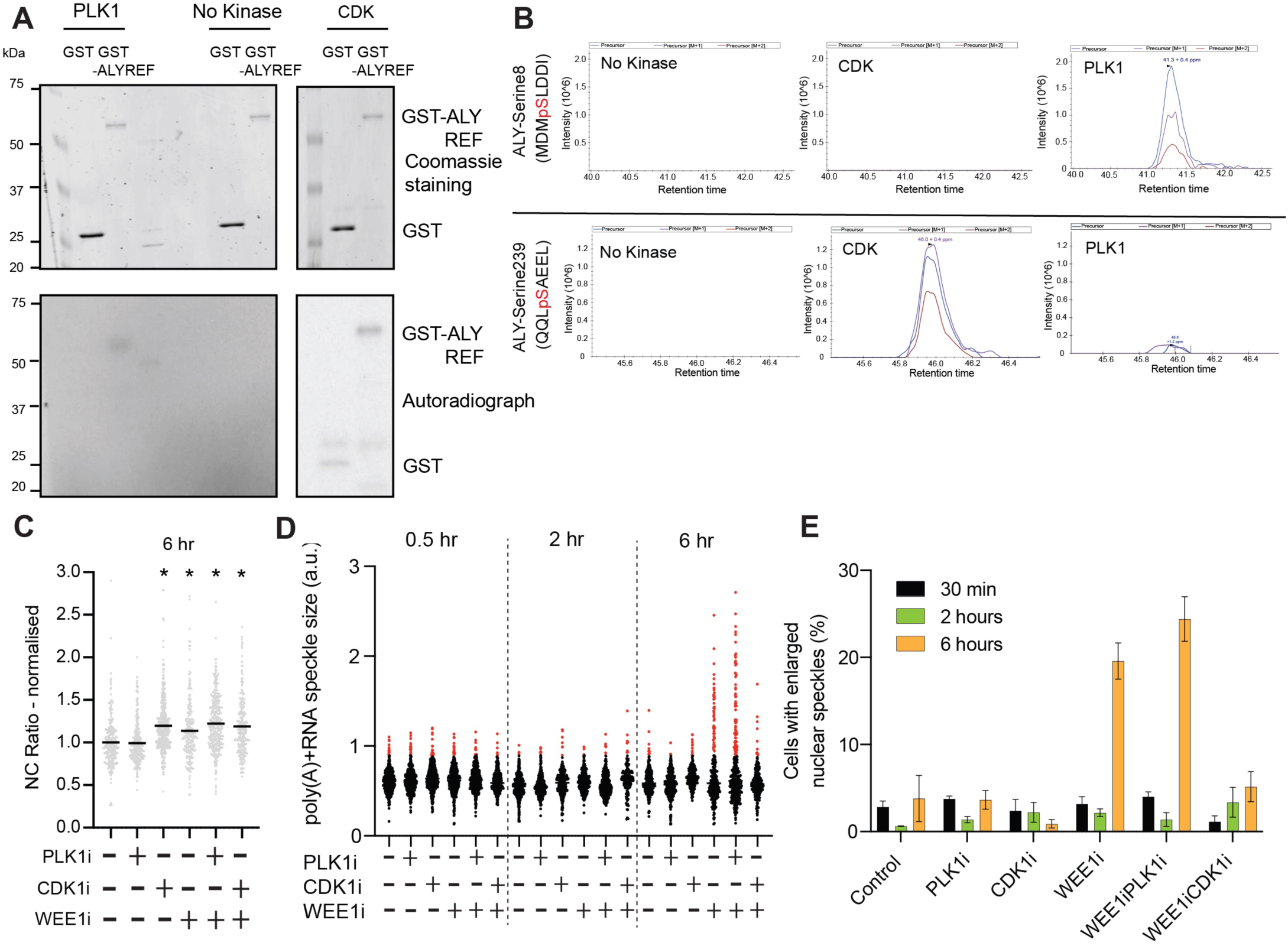
CDK1 signalling drives the nuclear accumulation of poly(A)+RNA to nuclear speckles in response to replication stress. **(A)** *In vitro* kinase assays with PLK1 or CDK1 and GST-ALYREF. GST alone was used as a negative control. Phosphorylation was assessed by autoradiograph. Coomassie staining of the indicated proteins are shown in the top panel. **(B**) PLK1 and CDK phosphorylate ALYREF at specific residues. Phospho-mass spec analysis of (**A**) was performed. Phosphorylated peptides that were identified are displayed in top and bottom panels. These were the only peptides that showed any phosphorylation. (**C**) N/C ratio of poly(A)+RNA following 6 hours of treatment with the indicated inhibitors. Cells were synchronised in S phase via double thymidine block and released before treatment. A minimum of 50 individual cells were analysed per experimental repeat (n=3). **(D**) Co-inhibition of WEE1 with CDK1 rescues the enlarged nuclear speckle defect. Poly(A)+ RNA speckle size of cells following the indicated kinase inhibition for 0.5, 2 and 6 hours. Cells with nuclear speckles greater than one standard deviation above the control cells are highlighted in red. **(E**) Cells with nuclear speckles greater than one standard deviation above the control from (**D**) expressed as a percentage. Graph represents data from 3 independent experiments with error bars showing SEM.

### CDK1 signalling drives the nuclear accumulation of poly(A)+RNA to nuclear speckles in response to replication stress

To determine whether CDK1 and PLK1 play a role in regulation of mRNA export during replication stress resulting from WEE1 inhibition, we assessed the localisation of poly(A)+RNA following WEE1 inhibition, WEE1 and CDK1 inhibition or WEE1 and PLK1 inhibition using chemical inhibitors specific for each kinase. CDK1 inhibition alone resulted in an mRNA export defect, but enlarged poly(A)+RNA speckles were not present (Figure 5C,D). Co-inhibition of CDK1 and WEE1, but not PLK1 and WEE1 reduced both poly(A)+RNA speckle size (Figure 5D) and the percentage of cells with enlarged nuclear speckles (Figure 5E) back to basal levels. Notably, these cells still have an mRNA export defect, similar to CDK1 inhibition alone (Figure 5C). Taken together, these findings implicate CDK1 signalling in driving the accumulation of poly(A)+RNA and ALYREF into nuclear speckles following replication stress.

### ALYREF and UAP56 co-operate to resolve R-loops and their interaction is regulated by phosphorylation

ALYREF and UAP56 are critical components of the TREX complex that interact directly with each other through the N and C-termini of ALYREF ^60–62^, and we identified these regions to be phosphorylated following replication stress (Figure 4C). To assess whether ALYREF phosphorylation might regulate UAP56 binding, we synthesised biotinylated peptides of ALYREF (residues 1-17 and residues 237-260) in unphosphorylated or phosphorylated forms. Using these peptides, we first determined whether UAP56 could be retrieved by ALY peptides from nuclear extract in a biotinylated pulldown assay. No interaction between UAP56 and either serine 239-phosphorylated or unmodified ALYREF (237-260) was detected, however we clearly observed a UAP56 interaction with ALYREF (1-17) (Figure 6A). Moreover, this interaction was abolished when ALYREF (1-17) was phosphorylated at serine 8 (Figure 6A), the residue phosphorylated by PLK1. Next, we measured interaction with recombinant UAP56 protein (Figure S6A). A similar selective interaction between UAP56 protein and ALYREF (1-17) in its unphosphorylated form was observed (Figure 6A). Previous studies in yeast and humans have suggested that ALYREF weakly promotes UAP56 ATPase activity ^63^ and UAP56 has been implicated in unwinding R-loops through its RNA-DNA helicase activity ^64^. As phosphorylation of ALYREF by PLK1 prevents its association with UAP56 (Figure 6A), we next determined whether ALYREF could stimulate the RNA:DNA hybrid unwinding activity of UAP56. To do this, we used an established R-loop mimic comprised of oligonucleotides. As previously demonstrated ^64^, we found that UAP56 is capable of removing RNA from this structure, but we also showed that recombinant GST-ALYREF, but not GST alone, stimulates this activity approximately 6-fold (Figure 6B,C). GST-ALYREF alone caused RNA partial displacement in an ATP-independent manner (Figure 6B,C). These findings are consistent with a recent report ^65^ and suggest that ALYREF can passively suppress R-loop formation by binding to displaced RNA and preventing annealing during strand “breathing”, but it can also greatly stimulate ATP-dependent active unwinding of R-loops by UAP56.

**Figure 6:**
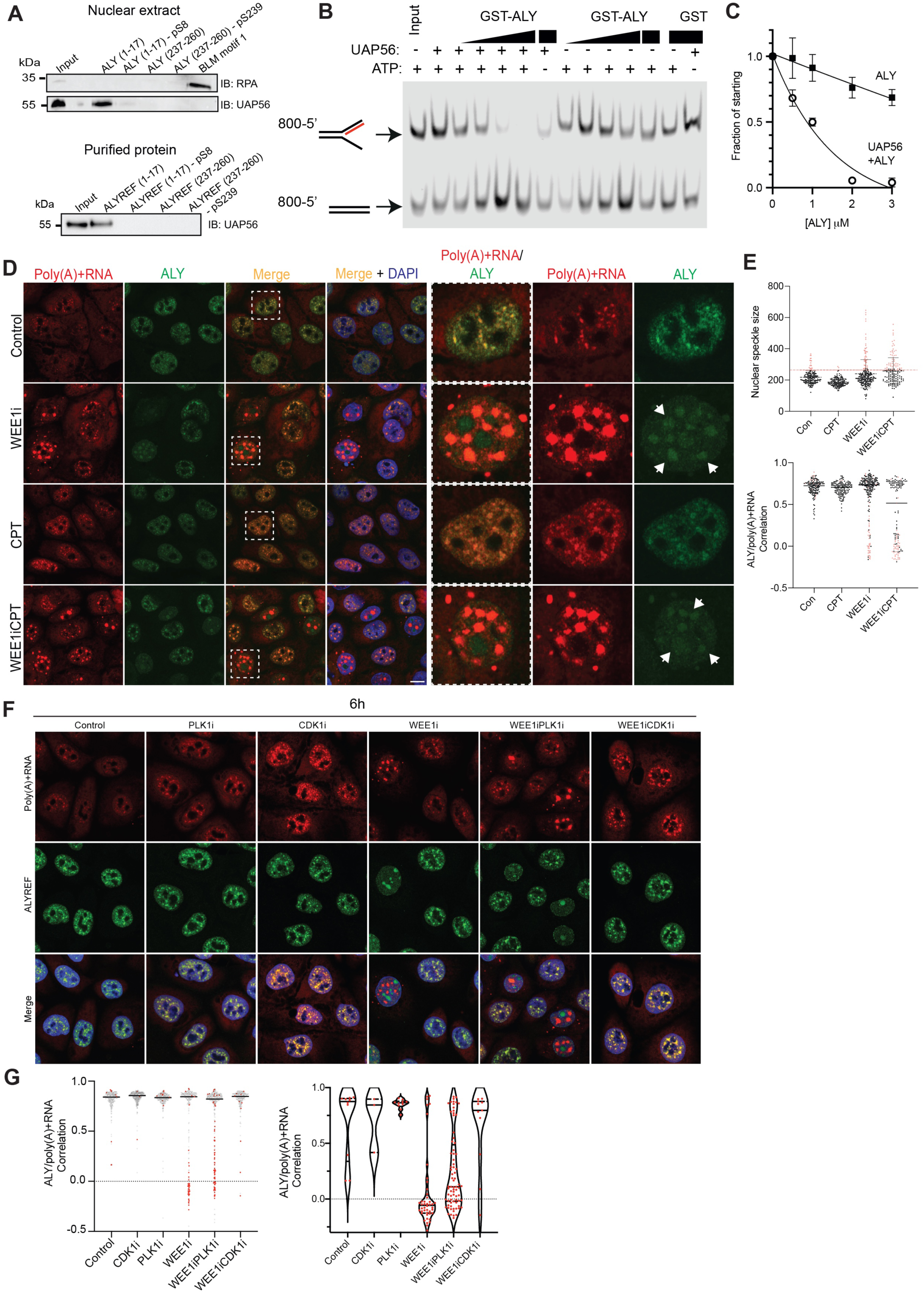
ALYREF and UAP56 co-operate to resolve R-loops, and their interaction is regulated by phosphorylation by CDK1. **(A**) Co-immunoprecipitation experiments with either nuclear extract (top panel) or purified UAP56 (bottom panel) and synthesised short ALYREF peptides containing phosphorylated and unphosphorylated S8 and S239 were performed. UAP56 levels were assessed by western blotting. Image is representative of 3 independent experiments. (**B**) *In vitro* RNA:DNA unwinding assays with purified UAP56 and increasing concentrations of GST-ALYREF with migratable 3′ and 5′ RNA:DNA flap structures. **(C**) Quantification of RNA:DNA hybrid resolution across 3 independent experiments, error bars represent standard deviation. (**D**) Confocal images of FISH for poly(A)+RNA with IF for ALYREF in cells treated with the indicated inhibitors, Images are representative of 3 independent experiments. Individual cells selected and displayed at a higher magnification are indicated by the white box. Cells with ALYREF staining in both nuclear speckles and nucleoli are shown and indicated by white arrowheads**. (E**) Nuclear speckle size and ALYREF/ poly(A)+RNA correlation from (**D**) with a minimum of 50 cells analysed per repeat for each treatment (n=3). Cells with nuclear speckles greater than one standard deviation above the control cells are highlighted in red. **(F**) Confocal images of FISH for poly(A)+RNA with IF for ALYREF in cells treated with the indicated inhibitors. Images are representative of 3 independent experiments. **(G**) Nuclear speckle size and ALYREF/ poly(A)+RNA correlation from (**F**) with a minimum of 50 cells analysed per repeat for each treatment (n=3) (left panel). Cells with nuclear speckles greater than one standard deviation above the control cells are highlighted in red. Cells with enlarged nuclear speckles were separated and their ALYREF/poly(A)+RNA correlation score is shown individually (right panel).

### mRNA export factors are mis-localised from enlarged nuclear speckles in cells with high levels of replication stress to prevent nuclear export of mRNA

Recent studies in 3D organisation of the nucleus have given rise to a new understanding of how nuclear compartments play a pivotal role in gene expression. However, while nuclear speckles are sub compartments known to contain high concentrations of poly(A)+RNA and RNA processing factors, including mRNA export factors, their exact role remains controversial ^23,24^. Our results thus far provide evidence that both ALYREF, and R-loops marked by phosphorylated RPA, are recruited to nuclear speckles during replication stress. We did not observe any obvious localisation of known R-loop resolution factors XPG or Senataxin ^66,67^ at nuclear speckles following replication stress (Figure S5B,C). This specific co-localisation may therefore be important for resolution by UAP56 of R-loops containing mature polyadenylated mRNA, in particular if they cannot otherwise be resolved at sites of replication stress. However, if this indeed the case, what happens if the R-loops are subsequently also not resolved by UAP56 at nuclear speckles?

A clue comes from our observation of wide-spread mis-localisation of mRNA export factors in cells with very large mRNA nuclear speckles. In these cells, ALYREF was mis-localised to nucleoli (Figures 6D,E and S5D), where it co-localised with nucleolar marker fibrillarin (Figure S5D). NXF1 and GANP were also mis-localised from the nuclear periphery in these cells (Figure S5E,F). This suggests that while these mRNAs have poly(A) tails, they are unable to be exported as there are no export factors in the vicinity to export them. Importantly, we observed cells with large mRNA nuclear speckles that contain ALYREF both in nuclear speckles and nucleoli (Figure 6D), suggesting that their relocalisation is a dynamic process. In support of this idea, we observed that in cells with CDK1 and WEE1 co-inhibition, large nuclear speckles containing poly(A)+RNA did not form, and ALYREF was not mis-localised to nucleoli (Figure 6F,G). Indeed, ALYREF phosphorylation on (CDK-phosphorylated) serine 239 is in a region (C-terminal) previously shown to be critical for its localization to mRNA nuclear speckles ^60^. In contrast, we found that in cells with PLK1 and WEE1 co-inhibition, large nuclear speckles containing poly(A)+RNA still formed. Furthermore, while we still observed ALYREF in nucleoli, a proportion of ALYREF remained in the enlarged nuclear speckles, as evidenced by the increase in ALYREF and poly(A)+RNA correlation in comparison to WEE1 inhibition alone (Figure 6G).

Our phosphoproteomics experiments revealed that PLK1 activity is temporally separated from CDK activation. Increased CDK activity occurs 2 hours post WEE1 inhibition, whereas PLK1 activity was only increased 6 hours post WEE1 inhibition (Figure 4). On ALYREF, phosphorylation at different residues was similarly temporally separated: serine 239 phosphorylation peaked at 2 hours post WEE1 inhibition, yet serine 8 phosphorylation was only detected 6 hours post WEE1 inhibition, when PLK1 activity is high (Figure 4C). These two results are consistent with our in vitro kinase data showing that PLK1 and CDK respectively phosphorylate serine 8 and 239 of recombinant ALYREF (Figure 5A-B). Importantly, it is phosphorylation of serine 8 that abolishes the interaction of the N-terminal ALYREF peptide with UAP56 (Figure 6A). Consistent with this observation, a dramatic relocalisation of UAP56 from nuclear speckles – such that ALYREF and UAP56 no longer co-localise and are in different cellular compartments (Figure S6B-F) – occurs in cells with enlarged nuclear speckles. These enlarged speckles are only observed in later timepoints and are accompanied by a reduction in the intensity of UAP56 in the nucleus (Figure S6E).

R-loops have been previously observed in the cytoplasm following Senataxin or BRCA1 depletion, or splicing inhibition ^42^. Interestingly, at early timepoints following replication stress and WEE1 inhibition, we observed the appearance of R-loops in the cytoplasm (Figure S6G-I). Using two orthogonal approaches, DNA/RNA immunoprecipitation and dot-blots from nuclear and cytoplasmic fractions using the S9.6 antibody, R-loops were detected in the nucleus and cytoplasm within an hour of treatment (Figure S6G-I). Together, these findings provide evidence that in cells with high levels of replication stress and increased nuclear speckle size, PLK1 phosphorylation of ALYREF blocks its interaction with UAP56, and disrupts the nuclear export of R-loop associated mature mRNA to the cytoplasm (Figure S7).

## Discussion

Our findings have revealed the existence of a multi-step cell cycle regulated checkpoint that exerts control of mRNA export in response to replication stress during S-phase. Based on our results we propose a model whereby TREX is recruited to sites of replication stress for R-loop resolution (Figure S7, left panel). When replication stress remains unresolved, CDK1 phosphorylation of ALYREF at serine 239 subsequently drives the relocalisation of poly(A)+RNA and R-loops to nuclear speckles for UAP56-mediated resolution (Figure S7, right panel). Then, if R-loops cannot be resolved by UAP56 and ALYREF at nuclear speckles, their association is abolished by a PLK mediated phosphorylation of ALYREF at serine 8. Dissociation of the two proteins promotes their re-localisation from nuclear speckles and prevents mRNA export (Figure S7, right panel). In this model, WEE1, CDK1 and PLK1 enforce a cell cycle checkpoint that is physical in nature, as mRNA export factors are mislocalised from the mRNA that they normally export to the cytoplasm. This checkpoint may serve to protect the cell from major sources of genome instability, by ensuring that R-loop associated mature mRNA is not exported to the cytoplasm.

During normal DNA replication in S-phase, this mechanism may function when cells undergo low levels of replication stress. Indeed, mRNA export factors are located in a prime position to resolve replication stress-associated R-loops because they are already physically associated with sites of ongoing transcription^9,10^, and they are further recruited to activated RPA during replication stress (Figure 2). UAP56 is the main helicase associated with TREX, it can interact with chromatin, and it possesses RNA-DNA helicase activity ^64^. Importantly, we show that the RNA/DNA hybrids accumulating in the absence of this mechanism contain polyadenylated RNA, suggesting that they are mostly processed, but not yet exported. If they cannot be resolved or released from sites of transcription, they are then recruited in a CDK dependent manner to nuclear speckles for resolution.

While the composition of nuclear speckles was first characterized over thirty years ago ^19–22^, the functional role of poly(A)+RNA in nuclear speckles remains contentious ^23,24^. Our results have unexpectedly uncovered an important functional role for nuclear speckles in processing of unresolved R-loops that contain poly(A)+RNA. As time progresses following replication stress, the nuclear speckles expand, and their mRNA content increases. This finding aligns with the emerging concept that speckles consist of a proteinaceous core enveloped by an RNA shell, whose size is correlated to RNA content ^68^. Strikingly, damaged single stranded DNA encapsulates the largest nuclear speckles after prolonged replication stress, as evidenced by co-localisation of phospho-RPA at their periphery. While these nuclear speckles contain mature polyadenylated mRNA, they do not contain dsDNA. But other recent observations suggest that a number of highly expressed active genes are also recruited to the periphery of nuclear speckles for transcription and processing ^9,24,29–33^. Indeed, the dramatic increase in the size of nuclear speckles that we observe following WEE1 inhibition and replication stress is due to an increase in poly(A)+RNA, providing further support to the idea that important cellular functions such as transcription, RNA processing ^24,29,30,69^ and in our case, R-loop resolution may take place in the RNA shell of the nuclear speckle.

mRNA export is extensively coupled to transcription and processing of mRNA throughout the cell cycle, and our unbiased screening approaches identify a major role for phosphorylation of ALYREF by cell cycle regulated kinases such as CDK1 and PLK1 in its regulation. It is also possible that phosphorylation of other mRNA export factors, in addition to ALYREF, may be a more widespread mechanism used by cells for regulating mRNA export. Given the recent identification of R-loops in the cytoplasm ^40–42^, it is tempting to speculate that an mRNA export checkpoint may function to retain R-loop associated mRNA in the nucleus under certain physiological circumstances. Indeed, we have been able to detect R-loops in the cytoplasm at early timepoints following replication stress. The physical separation of mRNA export factors from ostensibly export-competent mature polyadenylated RNA that we observe in large nuclear speckles when PLK1 activity is high, will prevent mRNA from being exported as there are no export factors in the vicinity to export them. Should this physical checkpoint fail, R-loop associated mRNA could be exported to the cytoplasm ^42^, emphasizing the importance of safeguarding regulation of mRNA export.

## Author contributions

VOW conceived the project. TDW performed the majority of experiments in this paper, with help from KTC, WBH and LHN. SVT, VM and AJD designed, performed and analysed the recombinant R-loop experiments, JL and EH performed and analysed chromatin mobility experiments. SJH performed and analysed Easyphos proteomic experiments. KJC, IN and KJS provided technical assistance with high-throughput screening and analysis experiments. KKB provided technical assistance. VOW supervised the study, performed and analysed experiments and wrote the paper.

**Declaration of interests**

VOW is a co-founder of exteRNA.

## Data and code availability

The mass spectrometry proteomics data have been deposited to the ProteomeXchange Consortium via the PRIDE partner repository with the dataset identifier PXD072996.

## Materials and Methods

### Cell culture

CAL51 breast adenocarcinoma cells were a gift from Professor Paul Edwards, Department of Pathology, University of Cambridge. They were cultured as monolayers in Dulbecco’s modified Eagles medium (Invitrogen) supplemented with 1 % (v/v) penicillin-streptomycin and 10 % (v/v) Foetal calf serum at 37°C in a 5 % CO_2_ incubator and passaged when 80-90% confluent. Cells were routinely tested for mycoplasma contamination.

### Generation of cell lines

#### Stable Cas9-CAL51 cell line

Cas9 lentivirus was kindly provided by the Victorian Centre for Functional Genomics (VCFG), Peter MacCallum Cancer Centre, Melbourne, Australia. Stable Cas9-expressing CAL51 cell lines were generated via lentiviral transduction of the FUCas9Cherry plasmid (Addgene # 70182) at 0.5 multiplicity of infection containing 8 μg/mL sequa-brene. CAL51 cells with high levels of Cas9-mCherry expression (∼top 20%) were sorted via FACS following preliminary experiments suggesting high expressing cells permitted more uniform editing efficiency. These cells maintain a high expression of Cas9 for upwards of 8 passages. Multiple aliquots of early passage Cas9 high CAL51 cells were frozen, thawed and passaged at least once the week before use, thus ensuring that cells of the same levels of Cas9 expression were used for all experiments.

#### Generation of pLIX_403-emGFP-ALYREF

pLIX_403-emGFP-ALYREF was generated by gateway cloning FLAG-emGFP-ALYREF (a gift from Professor Niels Gehring, Institute for Genetics, University of Cologne) into pLIX_403 (a gift from David Root, Addgene plasmid #41395) using the following primers: attB1GFP-ALYREF forward 5’: - GGGGACAAGTTTGTACAAAAAAGCAGGC TATGGACTACAAGGACGACGAT – 3’, attB2GFP-ALYREF forward 5’ - GGGGACCACTTTGTACAAGAAAGCTGGGTCTTAACTGGTGTCCATTCTCGC – 3’. Lentiviral particles were generated by co-transfecting 293T cells with pLIX_403-emGFP-ALYREF or pLIX_403-emGFP-ALYREF-S8A/S239A, pCMV-VSV-G (a gift from Bob Weinberg, Addgene plasmid #8454) and psPAX2 (a gift from Didier Trono, Addgene plasmid #12260). Viral supernatant was collected 48 hours after transfection, filtered through a 0.45-mm filter, and added to target cells in the presence of 10 µg/mLpolybrene (TR-1003-G, Merck). CAL51 cells with high levels of pLIX_403-emGFP-ALYREF expression were collected via fluorescence-activated cell sorting (FACS) following induction of emGFP-ALYREF expression with 1 µg/mL doxycycline. Cells were passaged in the presence of the selection antibiotic puromycin (1 µg/mL).

The following expression and mutagenesis plasmids were synthesized/purchased from Genscript Biotech: S-A Mutagenesis: pLIX_403-emGFP-S8A/S239A-ALYREF.

#### CAL51-TET-hRNAseH1::eGFP cells

Human RNase1 (ENST00000315212.4) with a nuclear localisation signal (***PKKKRKV***) substituting the region coding for the first 27 amino acids (RNAseH1 m27) which promotes nuclear accumulation and reduces mitochondrial targeting of RNAseH1 was synthesised as a gBlock fragment by IDT DNA and fused to monomeric EGFP by overlap extension using the PCR primers:

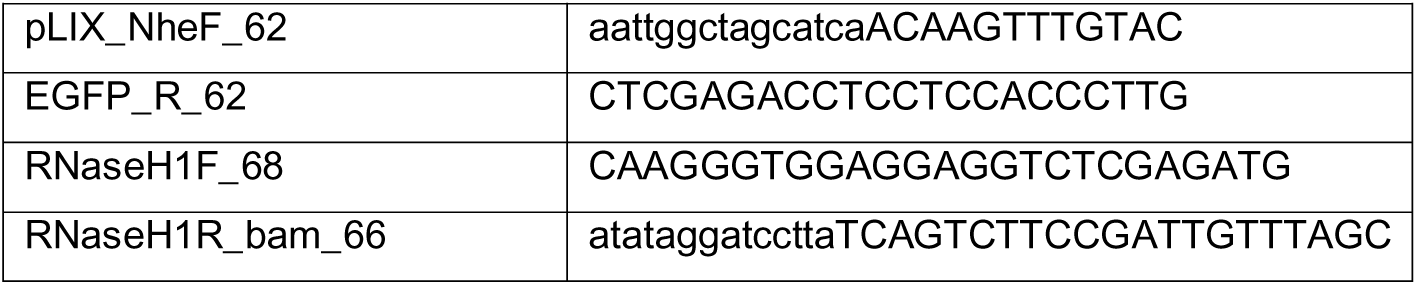

This cassette was cloned into the BamHI/NheI site of pLIX_403 (Addgene plasmid #41395) and verified by sanger sequencing. To generate the piggyBAC dual plasmids TET-inducible system PB-TET, the polyadenylation signal from the Bovine Growth Hormone (Addgene plasmid #121119) was fused to the RNAseH1::eGFP cassette by overlap extension using the PCR primers:

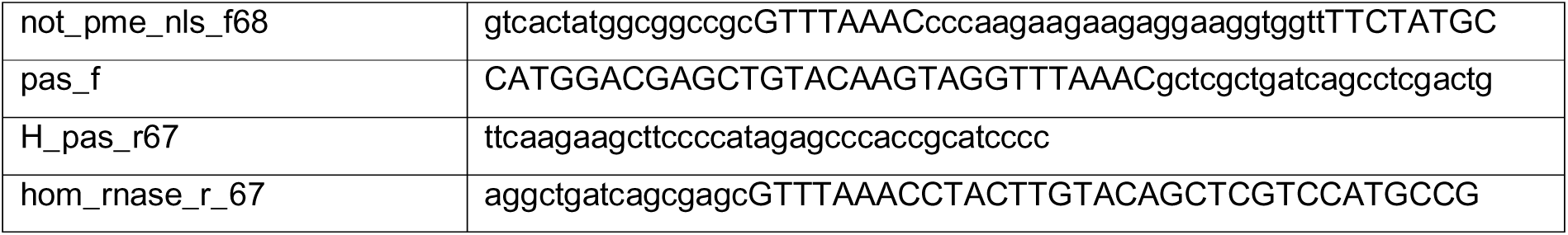

and cloned between the NotI and HindIII sites of PB-TET (Addgene plasmid #20909) and verified by sanger sequencing. Next the PGK-BSD resistance cassette (Addgene plasmid #26655) was fused to the SV40 polyadenylation signal (Addgene plasmid #41395) and cloned in the reverse orientation between the EcoRI sites of PB-TET with Gibson assembly using the PCR primers:

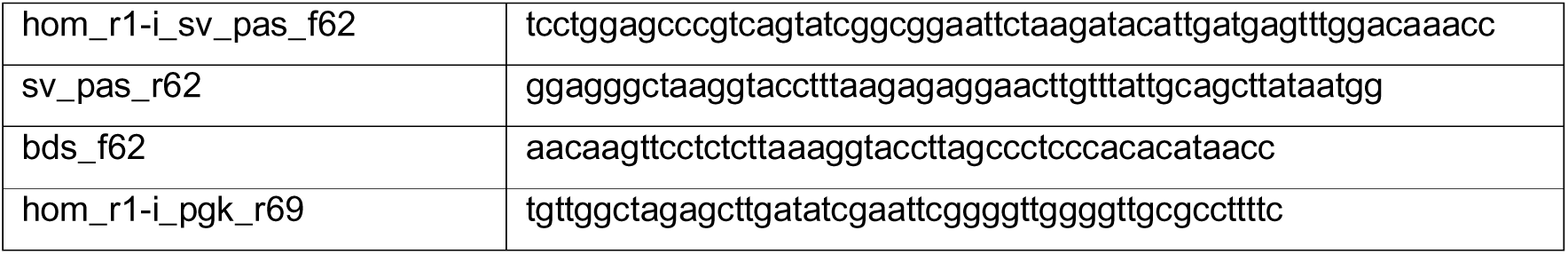

The final construct was verified by nanopore sequencing. The separate PB-EF1-rtTA-P2A-Hph plasmid was generated by successive rounds of digestion and self-ligation with XhoI and BamHI of the Addgene plasmid #122267 to remove the sequence upstream of the EF1-rtTA-P2A-Hph cassette but retain the PB-inverted repeats, the final plasmid was verified by restriction fragmentation.

CAL51-TET-hRNAseH1::eGFP cells were made by co-transfecting CAL51 cells with the activator and responder plasmids described above alongside pRM1024_CMV_PBase (a kind gift from Dr. Dane Vassiliadis) at a ratio 1:2:0.5, respectively using lipofectamine LTX. Cells were dual selected with 200 μg/mL hygromycin B and 5 μg/mL blasticidin (both from Sigma). Induction was performed by adding doxycycline hyclate (Sigma) to a concentration of 1 μg/ml.

### High content CRISPR-Cas9 and siRNA screens

CAL51-Cas9 cells or parental CAL51 cells (for RNAi) were screened in 384 well format using high content imaging for localization of polyA+RNA. On Day 1 of the experiment, crRNA or siGENOME SMARTpool RNAi library kinome (version 2009, Dharmacon RNAi Technologies, Horizon Discovery), as well as control plates prepared in house, were thawed at room temperature for one hour, pulse-spun at 60g, and then placed on ice. Subsequently, cells were trypsinized, counted, and diluted to give a final concentration of 1200 cells/well. DharmaFECT 1(DF1) was diluted in opti-MEM, and 25 µL of 2x DF1 lipid (1:500) was dispensed and mixed in master Plate A. Pre-complexed crRNA/tracrRNA from library plates were then dispensed into Plate A using a SciClone ALH3000 Lab Automation Liquid Handler (Perkin Elmer), followed by mixing and a 20-minute incubation at room temperature (RT). Half the master mix was then transferred to the B plate and cells were dispensed into the A and B plates using a BioTek EL406 (Agilent). Plates were incubated in a LiCONiCs incubator at 37°C with 5 % CO_2_ and at 24 hours post transfection, media was changed on all assay plates to address any potential toxicity from residual transfection lipid. Following a total of 72 hours transfection, using the BioTek EL406, cells were washed briefly with PBS, fixed in 4 % PFA for 5 minutes, washed with 1x PBS and incubated overnight in 100 % ethanol at −20°C. The next day, a high-throughput 384 well Cy5-oligod(T) FISH was performed using the BioTek EL406. Briefly, cells were washed twice with PBS, permeabilized for 10 minutes in 0.5% Triton X-100, washed with PBS for 5 minutes and incubated in 25 µL pre-hybridization buffer (2 x SSC, 20 % formamide, 0.2 % BSA, 1 mg/mL yeast tRNA in ultrapure water) for 30 minutes at 37°C. Following prehybridization cells were incubated in 25 µL hybridization buffer (prehybridization buffer + Cy5-oligo d(T) probe) for 3 hours at 37°C in a dark humidified chamber. After hybridization, cells were washed for 5 minutes twice with pre-warmed 2 x SSC and 20 % formamide followed by 2 x SSC at 42°C. Cells were then washed twice with 1 x SSC for 5 minutes each at RT followed by 1 x PBS twice for 5 minutes at RT. The first PBS wash contained DAPI (0.2 µg/mL). All liquid dispensing was performed by a BioTek EL406, wash volumes were 50µL and liquid was flicked out of plates in between steps to ensure removal of any residual dead volume.

Images were captured using the Cellomics ArrayScan VTI automated microscope at 20 x magnification (2 x 2 binning), 25 fields/well in widefield. The following channels were used: (1) DAPI (385 nm) and (2) Cy5 (650 nm). The in-built Compartmental Analysis BioApplication was used to perform automated image analysis at the time of acquisition. The DAPI channel was first used to identify primary objects (nuclei) using the Fixed intensity thresholding method. The secondary objects (cytoplasm) were then identified by creating a 7 pixel-wide ring around the primary object mask. The nuclear – cytoplasm intensity (N/C) ratio was calculated for each cell by dividing the nucleus mean intensity by the cytoplasm mean intensity in the Cy5 channel. Where possible, two thousand cells per well were measured. The single cell values were then aggregated into mean per well values. Robust Z scores were calculated and used for hit identification. Gene targets with N/C ratio Z scores >2 were selected as hits for further investigation.

### siRNA-mediated depletion

Depletions of different proteins were induced by reverse transfection using small interfering RNA and DF1 transfection reagent (Dharmacon) as previously described (Wickramasinghe et al., 2013). Briefly, Cells were transiently transfected with the indicated small interfering RNAs (siRNAs) using lipid-based transfection reagent DF1. The appropriate volume (1:1000 DF1 final) of lipid was added to Opti-MEM (Invitrogen) and incubated at RT for 5 minutes. In the meantime, siRNAs were diluted in Opti-MEM to provide a final concentration of 25 nM. siRNA/DF1 were then combined and allowed to complex at RT for 20 minutes. Transfection mixtures were added to wells followed by seeding of cells and incubation in 5 % CO_2_ at 37°C for 24 hours before the medium was replaced with fresh growth medium. Transfected cells were incubated in 5 % CO_2_ at 37°C for 48 or 72 hours. Seeded cell density was selected according to the timepoint, with 2.5 × 10^5^ cells seeded for 72 hour and 0.5 × 10^5^ cells seeded for 48 hour timepoints 9 per well in a 6 well plate). Efficiency of siRNA-mediated depletion at the protein level was examined by sodium dodecyl sulphate polyacrylamide gel electrophoresis (SDS-PAGE) and western blotting with the appropriate antibodies.

### crRNA-mediated depletion

The indicated guides were reconstituted to 20 μM in 10 mM Tris–HCl buffer (10 mM Tris, 2M HCl). Guides for individual genes were pooled prior to transfection. Pooled guides and non-targeting guide AAVS1 were complexed with Edit-R tracrRNA at a ratio of 1:1 for 20 minutes with DF1 transfection reagent in Opti-MEM (1:1000 final) and transfected at a final concentration of 25 nM into 2 mL of CAL51-Cas9 cells in a 6 well plate. Seeded cell density was selected according to the timepoint, with 2.5 × 10^5^ cells seeded for 72 hour and 0.5 × 10^5^ cells seeded for 48 hour timepoints. Cells were incubated in 5 % CO_2_ at 37°C for 24 hours when the medium was replaced with fresh growth medium. Transfected cells were incubated for the indicated periods of time before harvesting. Efficiency of CRISPR-mediated depletion of target proteins was examined by SDS-PAGE and western blotting with the appropriate antibodies.

### Synchronisation of cells in S-phase by double thymidine block

8 × 10^5^ cells were plated in media supplemented with 2 mM filter-sterilised thymidine in a 6 well plate at 1700 and incubated overnight. The following morning at 0900, cells were washed once with warm PBS and then incubated in cell culture medium for 8 hours precisely. 2 mM filter-sterilised thymidine was then added at 1700. The following morning at 0900 cells were released or held at G_1_/S and subsequently cultured in 5 % CO_2_ at 37°C in the presence of the indicated chemical compounds for the indicated periods of time. Synchronisation of cells at the G_1_/S boundary was assessed by DNA content analysis by flow cytometry.

### Drug treatment of synchronised cells

Cell culture medium was aspirated and replaced by the appropriate volume of drug-containing medium. Final concentrations of drugs were: WEE1i: 500nM, CDK1i: 9µM, CPT: 10µM. Cells were incubated at 37°C in a 5 % CO_2_ incubator for the indicated periods of time before harvesting.

### Cell cycle and DNA content analysis by flow cytometry

Trypsinised cells were resuspended in hypertonic solution containing 0.5 mM sodium citrate, 1% Triton X-100, 25 µM propidium iodide in ultrapure water. Cells were incubated overnight at 4°C in the dark before being resuspended in 1 % FCS in PBS and analysed using flow cytometry for PI fluorescence in the FL2-A (linear) and FL3-A (log) channels. Analysis of cell cycle phases of remaining viable cells was conducted by gating on viable cells (thereby excluding the sub-G_1_ fraction) in FL2-A data and placing markers for G_1_, S and G_2_ cell cycle phases. All flow cytometry data was acquired on FACS Canto II flow cytometers (BD Biosciences). Gates were set around single cells (FSC-Height vs FSC-Area) and debris were excluded by gating morphology (FSC-Area vs SSC-Area). At least 1×10^4^ cells were analysed for each sample. All flow cytometry data analyses were performed using FlowJo software.

### Protein analysis

Adherent cells were trypsinised, collected and wells rinsed with PBS. Cells were counted, centrifuged at 500 g for 5 minutes and washed with ice cold PBS. Cells were lysed with LDS lysis buffer (nuPAGE) containing 20 mM dithiothreitol (DTT) and boiled at 95°C for 10 minutes. Extracted protein was frozen at −80° C for analysis by SDS-PAGE and western blotting.

### Fluorescent in situ hybridisation (FISH) and FISH with Immunofluorescence

Cells seeded onto coverslips were washed with 1 x PBS for 5 minutes at RT, prior to fixation with 4 % paraformaldehyde (PFA) for 5 minutes at RT. Coverslips were then washed with 1 x PBS for 5 minutes before placing in 100% ice cold ethanol overnight (or longer until staining is performed). Fixed cells were permeabilised with 0.5% Triton X-100 in PBS for 10 minutes at RT, washed once with PBS at RT and incubated in pre warmed pre-hybridisation buffer (2 x SSC, 20 % formamide, 0.2 % BSA, 1 mg/mL yeast tRNA in ultrapure water) for 15 minutes at 37°C. For FISH-IF, coverslips were incubated in pre-hybridisation buffer for 30 minutes. Following pre-hybridisation, cells were incubated with hybridisation buffer (pre hyb buffer + cy3-oligo d(T) probe and 10% dextran sulphate) for 3 hours at 37°C in the dark in a humidified chamber before being washed for 5 minutes twice with pre warmed 2 x SSC plus 20 % formamide followed by 2 x SSC at 42°C. Cells were then washed twice with 1 x SSC for 5 minutes each at RT followed by PBS twice for 5 minutes at RT. The first PBS wash contained DAPI 0.2 µg/mL. For FISH-IF, antibodies for the indicated protein were included in the hybridisation buffer. For FISH-IF, the first 1 x SSC wash contained the appropriate secondary antibody diluted at 1:500 for 30 minutes at 37°C, with subsequent wash steps the same as the above regime. Following washes, coverslips were mounted with DAKO mounting medium on microscopy slides and stored at 4° C. Mounted coverslips were warmed to RT before imaging.

### Immunofluorescence

Cells seeded on coverslips were washed with 500 μL PBS for 5 minutes at RT. Cells were then fixed in 500 μL 4 % PFA for 5 minutes at RT, washed with 500 μL PBS for 5 minutes before being left in 1 mL of 100 % ice cold ethanol overnight (or longer). Fixed cells were then permeabilised with 500 μL permeabilisation buffer (0.1 % Triton X-100, 0.02 % SDS in PBS) for 10 minutes at RT. Following permeabilization, cells were blocked for 30 minutes with 500 μL blocking buffer (0.1 % Triton X-100, 0.02 % SDS, 1 % BSA in PBS) and incubated with 60 μL of the indicated primary antibody (in blocking buffer) for 1 hour at 37°C in a dark humidified chamber. Cells were then washed 3 x 5 minutes each in 500 μL blocking buffer at RT. Secondary antibodies were diluted in blocking buffer and 60 μL added dropwise to form a meniscus on top of the coverslip before incubation for 1 hour at 37°C in the dark in a humidified chamber. Following incubation, cells were washed 3 times with 500 μL blocking buffer with DAPI added at a dilution of 1:5000 in the second wash. Following washes, coverslips were mounted with DAKO mounting medium on microscopy slides and stored at 4° C. Mounted coverslips were warmed to RT before imaging.

### Cell-cycle analysis using Click-iT EdU

CAL51 cells grown on coverslips were spiked with 10 μM of EdU for 10 minutes to label S phase cells. Cells were then washed with 500 μL of 1x PBS for 5 minutes at RT, fixed with 500 μL 4 % PFA for 5 minutes at RT, washed with 500 μL 1 x PBS for 5 minutes and finally placed in 1 mL 100 % ice cold ethanol overnight (or longer). Fixed cells were permeabilised with 500 µL of 0.5% Triton X-100 in PBS for 10 minutes at RT and washed once with 500 µL of 1 x PBS at RT. Following permeabilisation, cells were washed twice with 500 µL of 3 % BSA for 5 minutes each and coupling of EdU to the Alexa fluor substrate occurred in the dark in the Click-iT reaction mixture (Thermo Fisher Scientific), prepared according to manufacturer’s instructions.

### Confocal laser scanning microscopy and FLIM data acquisition

Fixed cell fluorescence intensity and lifetime microscopy measurements were performed on an Olympus FV3000 laser scanning microscope coupled to a 488 nm pulsed laser operated at 80 MHz and an ISS A320 Fast FLIM box. A 60x water immersion objective 1.2 NA was used for all experiments and the cells were imaged at room temperature. First, three colour intensity images (512 x 512-pixel frame size, 12.5 µs/pixel, 45 nm/pixel) were acquired of each selected nucleus to: (1) verify the presence of the histone FRET pair (H2B-eGFP and H2B-mCh) with an acceptor: donor ratio > 1, and (2) record the spatial localisation of AF647 marked Wee1iCPT. This involved sequential imaging of a two-phase light path in the Olympus FluoView software. The first phase was set up to image H2B-eGFP and H2B-mCh via use of solid-state laser diodes operating at 488 and 561 nm, respectively, with the resulting signal being directed through a 405/488/561/640 dichroic mirror to two internal GaAsP photomultiplier detectors set to collect 500–550 nm and 600–650 nm. The second phase was set up to image AF647 via use of a solid-state laser diode operating at 640 nm with the resulting signal being directed through a 405/488/561/640 dichroic mirror to an internal GaAsP photomultiplier detector set to collect 650–700 nm. Then in each nucleus selected, a FLIM map of H2B-eGFP was imaged within the same field of view (256 × 256-pixel frame size, 20 µs/pixel, 90 nm/pixel, 20 frame integration) using the ISS VistaVision software. This involved excitation of H2B-eGFP with the external pulsed 488 nm laser (80 MHz) and the resulting signal being directed through a 405/488/561/640 dichroic mirror to an external photomultiplier detector (H7422P-40 of Hamamatsu) that was fitted with a 520/50 nm bandwidth filter. The donor signal in each pixel was then subsequently processed by the ISS A320 FastFLIM box data acquisition card to report the fluorescence lifetime of H2B-eGFP. All FLIM data were pre-calibrated against fluorescein at pH 9 which has a single exponential lifetime of 4.04 ns.

### Phasor FLIM analysis of histone FRET

The fluorescence decay recorded in each pixel of an acquired FLIM image was quantified by the phasor approach to lifetime analysis. As described previously ^70–72^, this results in each pixel of a FLIM image giving rise to a single point (phasor) in the phasor plot, which when used in the reciprocal mode enables each point in the phasor plot to be mapped to each pixel of the FLIM image. Since phasors follow simple vector algebra, it is possible to determine the fractional contribution of two or more independent molecular species coexisting in the same pixel. For example, in the case of two independent species, all possible weightings give a phasor distribution along a linear trajectory that joins the phasors of the individual species in pure form. While in the case of a FRET experiment, where the lifetime of the donor molecule is changed upon interaction with an acceptor molecule, the realization of all possible phasors quenched with different efficiencies describes a curved FRET trajectory in the phasor plot that follows the classical definition of FRET efficiency ^71,73^. In the context of the histone FRET experiments presented, the phasor coordinates of the unquenched donor (H2B-eGFP) and background (cellular autofluorescence) were first determined independently in fixed cells transfected versus un-transfected with H2B-eGFP. This enabled definition of a baseline from which a FRET trajectory could be extrapolated and then used to determine the dynamic range of FRET efficiencies that describe chromatin nanostructure in the cell system ^73,74^. From superimposition of this FRET trajectory with the combined phasor distribution measured for H2B-eGFP in fixed cells co-transfected with H2B-mCh, we find a shift in the H2B-eGFP donor lifetime from approximately 2.5 ns to 2.0 ns. We therefore defined two cursors centred at these phasor coordinates to spatially map where chromatin is open (blue cursor) versus compact (orange cursor) throughout a FLIM data acquisition in a fixed nucleus and quantify overall chromatin network compaction based on the fraction of pixels exhibiting histone FRET ^73,74^. All FLIM-FRET quantification was performed in the SimFCS software developed at the Laboratory for Fluorescence Dynamics and a Matlab code specifically designed to mask FLIM data according to an IF signal ^74^, which enabled calculation of the chromatin nanostructure defined by WEE1i + CPT localisations.

### Structure bound protein co-immunoprecipitation

CAL51 cells were seeded in 10cm dishes at 5 × 10^6^ cells overnight. The following day, cells were washed with ice cold PBS, scraped in 10 mL of digitonin buffer (50 µg/ml digitonin, 100 mM NaCl, 10 mM Tris, 1x Roche Protease inhibitor cocktail (10 µL per mL), RNase out (1.5 µL per mL)). and incubated for 15 minutes on ice. Cell nuclei were pelleted by centrifugation for 3 minutes at 1000 x g at 4°C, then washed with wash buffer (100 mM NaCl, 10 mM Tris, pH 8.0, 1 x Protease Inhibitor cocktail (Roche)), and lysed with 400 μL of low salt lysis buffer (150 mM NaCl, 10 mM Tris pH 8.0, 1x Protease Inhibitor cocktail (Roche)) for 5 minutes on ice, pelleted at 15 000 g for 5 minutes, the nuclear soluble fraction was then precipitated in 6 volumes of ice-cold acetone and stored at −20° C for later processing. The pellet containing structure bound proteins was then extracted in high salt buffer (10 mM Tris pH 8.0, 500 mM NaCl, 0.2% NP-40 and 1x Protease Inhibitor cocktail (Roche)) for 15 minutes on ice and centrifuged at max speed for 30 minutes at 4° C. Nuclear lysate was diluted in 800 μL lysis buffer without NaCl per reaction and pre-cleared with 60 μL of Protein G dynabeads (Thermo Fisher Scientific) for 30 minutes at 4°C with gentle rotation. 5 μg of either anti RPA2 antibody (Abcam) or mouse IgG isotype control (Thermo Fisher Scientific) was then added to nuclear lysates and incubated for 2 hours at 4°C with gentle rotation. Samples were then incubated with 60 μL Protein G dynabeads (Thermo Fisher Scientific) overnight at 4°C with rotation. Finally, beads were washed 3 times with wash buffer and proteins were eluted in 60 μL 1X LDS sample buffer (Invitrogen) with 50 mM DTT and boiled for 10 minutes at 95° C. Effective separation of nuclear-soluble proteins from structure-bound proteins was confirmed by SDS-PAGE and western blotting for nuclear-soluble Exportin-5 (XPO5) and structure-bound Histone H3.

### DNA:RNA immuno-precipitation (DRIP) – qPCR

1 × 10^7^ CAL51 cells were treated with the indicated drug for 8 h prior to fixation. Crosslinking of cells was performed with 1% PFA for 10 minutes, then quenched with 500 mM glycine (pH 6) for 5 minutes at RT. Cell lysis and DNA extraction was performed using a NucleoSpin Tissue kit (Machery Nagel) following a modified cell lysis protocol. Briefly, cells were lysed in 800µL T1 + B3 lysis buffer (1:1) supplemented with 25 µL Proteinase K and incubated at 37°C with mild shaking overnight. The following day, DNA purification was performed as per manufacturers protocol and nucleic acids were eluted in two rounds of 100 µL elution buffer warmed up to 65° C. 12-14 µg DNA was then sonicated in a total volume of 300 µL of Tris buffer (10 mM Tris-HCl (pH 8.5), 300 mM NaCl) for 13 x 30 sec (30 sec ON, 30 sec OFF, 20% power) using a probe sonicator to yield an average DNA fragment size of ∼300 bp. Magnetic Dynabeads Protein A (ThermoFisher Scientific) were pre-blocked for 1 hour with 0.5 % BSA in PBS/EDTA and incubated with 10 µg of S9.6 antibody in 200 µL IP/lysis buffer (50 mM Hepes/KOH at pH 7,5; 0.14 M NaCl; 5 mM EDTA; 1% Triton X-100; 0,1 % Na-Deoxycholate) at 4°C for 4 hours with rotation. For RNAse H treated samples, 2 µL RNAse H enzyme (New England Biolabs) in NEB buffer was added to 6 µg of digested genomic DNA overnight at 37°C with mild shaking. The following day a further 1 µL RNase H enzyme was added for 1h. 6 µg of digested genomic DNA was the added to the S9.6 bound beads mixture and gently rotated at 4° C, overnight. Beads were recovered and washed 2 times each with 1 mL of low salt IP/lysis buffer (50 mM Hepes/KOH pH 7.5, 0.14 M NaCl, 5 mM EDTA pH 8, 1 % Triton X-100, 0.1 % Na-Deoxycholate), 1 mL of lysis buffer (high salt, 50 mM Hepes/KOH pH 7.5, 0.5 M NaCl, 5 mM EDTA pH 8, 1 % Triton X-100, 0.1 % Na-Deoxycholate), 1 mL of wash buffer (10 mM Tris-HCl pH 8, 0.25 M LiCl, 0.5 % NP-40, 0.5 % Na-Deoxycholate, 1 mM EDTA pH 8) and 1 mL of TE (100 mM Tris-HCl pH 8, 10 mM EDTA pH 8) at 4° C. Elution was performed in 100 µL of elution buffer (50 mM Tris-HCl pH 8, 10 mM EDTA, 1 % SDS) for 15 minutes at 65° C. After purification by NucleoSpin Gel and PCR Clean-up Kit (Macherey-Nagel), the recovered DNA was analyzed by quantitative real-time PCR (qPCR) using LightCycler 480 SYBR Green I Master Mix (Roche) on Roche LightCycler 480 platform according to manufacturer’s instructions.

### Nuclear and cytoplasmic DRIP – qPCR and dot-blot

18 ×10^6^ CAL51 cells synchronised and treated with DMSO or WEE1 inhibitor and camptothecin for 1, 2 and 4 hours, were washed in ice-cold PBS and pelleted by centrifugation and fractionated in 9 ml of digitonin buffer (50 µg/ml digitonin (MP Biomedicals, 215948090), 100 mM NaCl, 10 mM Tris-HCl pH 8.0) for 10 minutes on ice. The cytoplasmic or nucleoplasmic fractions were recovered and incubated in 10 mM EDTA pH 8.0, 0.4 % SDS and 40 µg/mL of proteinase K (Thermo Fisher Scientific, 25530049) for 90 min at 37^0^C. Cytoplasmic and nucleoplasmic nucleic acids were extracted using Phenol:Chloroform:Isoamyl alcohol 25:24:1 and precipitated with 1:10 volume of 3 M sodium acetate pH 5.2 and 2.5 volumes of 100 % ethanol. Samples were resuspended in 100 µL RNase-free water. An equal volume of each fraction, which was equivalent to the same number of cells, from each treatment was treated with 0.4 U/µL of RNaseOUT (Invitrogen, 10777019) or RNase H (New England Biolab, M0297S) for 3 hours at 37^0^C. For dot-blot samples, nuclear nucleic acids from 0.9 × 10^5^ cells, and cytoplasmic nucleic acids from 2.7 × 10^6^ cells were prepared for 2-fold serial dilutions using RNase-free water and blotted on N+ Hybond membrane (Amersham). Membrane was then blocked with 5 % skim-milk in TBS-0.1% Tween-20 for 1 hour at room temperature, then incubated overnight with S9.6 antibody. Membrane was washed three times with TBS-0.1% Tween-20, incubated in secondary antibody conjugated to HRP for 1 hour at room temperature, then washed in TBS-0.1% Tween-20 before developing.

For DRIP-qPCR samples, nuclear fractions were then diluted to 300 µL volume with 300 mM NaCl and 10 mM Tris-HCl pH 8.0 for sonication with a probe sonication at 20 % amplitude for 2 minutes. 10 % of nuclear fraction was used for input, with DNA being extracted using Qiagen MinElute Reaction Cleanup kit (Qiagen, 28204). Meanwhile, 16 µg of S9.6 antibody was bound to 50 µL Protein G Dynabeads (Invitrogen) in 1 x binding buffer (20 mM Tris-HCl pH 8.0, 2 mM EDTA pH 8.0, 1 % Triton X-100, 150 mM NaCl, 0.5 % sodium deoxycholate) for 5-6 hours at 4^0^C. For immunoprecipitation, cytoplasmic and nucleoplasmic fractions were diluted to 600 µL final volume in 1x binding buffer, then mixed with S9.6 antibody-bound beads and incubated overnight on rotation at 4^0^C. Beads were washed twice each with TSE buffer (20 mM Tris-HCl pH 8.0, 2 mM EDTA, 1 % Triton X-100, 0.1 % SDS, 150 mM NaCl) and TE buffer. Nucleic acid was eluted in 200 µL elution buffer (50 mM Tris pH 8.0, 10 mM EDTA, 0.5 % SDS, 80 µg/ml proteinase K) for 60 min at 50^0^C. Eluted RNA was purified with Qiagen RNeasy MinElute kit (Qiagen, 74204). RNA and input DNA were then used for reverse transcription and real-time qPCR.

### Phosphoproteomics Methods

Phosphoproteomics experiments were performed with four biological replicates (*n* = 4). <u>Phosphopeptides</u> were enriched using the EasyPhos workflow as previously described ^56^. Briefly, cells were lysed in SDC buffer (4% Sodium <u>deoxycholate</u>, 100 mM Tris pH 8.5) and immediately heated for 5 minutes at 95° C. Lysates were cooled on ice, and sonicated with a tip-probe sonicator (50% output power, 30 s). An aliquot of lysate was taken and diluted 1:5 in 8 M Urea from which protein concentration was determined by BCA assay (Thermo Fisher Scientific). Aliquots corresponding to 250 µg of protein were subsequently diluted in SDC buffer into a 96-well deep-well plate, reduced and alkylated at 45° C for 5 minutes by the addition of 10 mM Tris (2-carboxyethyl)phosphine (TCEP)/40 mM 2-Chloroacetamide (CAA) pH 8, and digested by the addition of 1:100 Lys-C and Trypsin overnight at 37°C with agitation (1,500 rpm). After digestion, phosphopeptides were enriched in parallel according to the EasyPhos workflow as described ^56^. Eluted phosphopeptides were dried in a SpeedVac concentrator (Eppendorf) and resuspended in MS loading buffer (0.3% TFA/2% acetonitrile) prior to LC-MS/MS measurement.

### LC-MS/MS Measurement

Phosphopeptides were loaded onto a 60 cm column fabricated in-house with 75 μM inner diameter fused silica packed with 1.9 μM C18 ReproSil particles (Dr. Maisch GmBH), which was maintained at a constant temperature of 60° C using a column oven (Sonation). Peptides were separated with a U3000 HPLC system (Dionex, Thermo Fisher Scientific) connected to a Q Exactive HF X benchtop Orbitrap mass spectrometer (Thermo Fisher Scientific) using a NanoSpray Flex ion source (Thermo Fisher Scientific). A binary buffer system of 0.1% (v/v) formic acid (buffer A) and 80% (v/v) acetonitrile/0.1% (v/v) formic acid (buffer B) was used, and peptides were eluted at a flow rate of 350 nl/min with a gradient of 3 – 19% buffer B over 40 minutes, followed by 19 – 41% buffer B over 20 minutes for a total peptide elution duration of ∼1 hour. Peptides were analysed with a full scan (350 – 1,400 m/z; R = 60,000 at 200 m/z) at a target of 3e6 ions, followed by up to ten data-dependent MS2 scans using HCD (target 1e5; max. IT 50 ms; isolation window 1.6 m/z; NCE 27%; min. AGC target 2e4), detected in the Orbitrap mass analyser (R = 15,000 at 200 m/z). Dynamic exclusion (30 s) and Apex trigger (2 to 4 s) were enabled.

### MS data processing

RAW MS data was processed in the MaxQuant software environment (<u>Cox and</u> <u>Mann, 2008</u>) (version 1.6.0.9), searching against the Human UniProt Reference database (June 2019 release), using default settings with the addition of ‘Phospho(STY)’ as a variable modification and ‘Match between runs’ switched on for all analyses. Data analysis was performed using the Perseus software package (<u>Tyanova and Cox, 2018</u>).

### Protein Purification

Hi 5 cells (200ml at 2×10^6^ cells/ml) were infected with 5 ml of UAP56 P2 virus and grown for 3 days. The cells were harvested by centrifugation and lysed by sonication in 20 mM TEA pH 7.5/150 mM NaCl/10 % Glycerol/ 1 mM DTT/1 mM EDTA/1 x mPI (Sigma Aldrich). The lysate was clarified by centrifugation. 250 μl of M-2 flag resin (Sigma Aldrich) was added and 90 minutes after rotation at 4°C the resin was washed and eluted with 100 μg/ml 1x flag peptide. The flag pure fractions were pooled and diluted 1:1 with 40 mM TEA pH7.5/ 5 % glycerol/2 mM 2ME. This material was bound to a MonoQ (Cytvia) column and eluted with a gradient of NaCl from 50 mM to 500 mM (40 mM TEA pH 7.5/ NaCl/ 5% glycerol /2 mM 2ME). Pure UAP56 containing fractions were pooled and concentrated using a 30 MWCO (Thermo Fisher Scientific) filter 15ml unit. Final volume recovered was 330 μl at 1.8 mg/ml.

### UAP56 and ALYREF peptide Co-IP

Magnetic protein G beads (Invitrogen) were washed with PBS and incubated with 5 mM of the indicated short ALYREF peptide (see Table S2 for sequences) in PBS for 30 minutes at room temperature with shaking. Subsequently, the beads were washed 5 times with 0.1 % BSA and resuspended with 5 µg of UAP56 in binding buffer (50mM HEPES pH 7.5, 100 mM NaCl, 1 mM EDTA, 1 mM DTT, 0.5% Triton X-100, 10% glycerol in water) and incubated for 2 hours at 4°C on a rotator. The beads were then washed 4 times with binding buffer before protein was eluted with 1x LDS buffer supplemented with 50 mM DTT, at 95°C for 10 minutes. Levels of UAP56 were assessed by western blot.

### R-loop generation

R-loops were generated with annealed oligonucleotide in a two-step reaction as previously described ^75^. Briefly, the steps are Step 1 set up 2 separate annealing reactions: 1) 4µL each 100 μM oligos A+C 2) 4µLeach 100 μM oligos B+L in 2 μl 5 x annealing buffer (25 mM Tris-HCL pH 7.5, 5 mM MgCl_2_, 50 mM NaCl). In a PCR machine, both tubes were heated at 98°C for 5 min, then cooled at a rate of 1°C/minute until they reached 4°C. Once cooled, tube 1 was mixed with tube 2 and incubated at 37°C for 30 mins, followed by 30 mins at room temperature. R-loops were gel purified, the band was excised and eluted in 400 μl of TMgN buffer (10 mM Tris pH 7.4, 5 mM MgCl_2_, 50 mM NaCl).

A: 5’ACGCTGCCGAATTCTACCAGTGCCTTGCTAGGACATCTTTGCCCACCTGCAGGTTCACCC3’

C: RNA-5’GGGUGAACCUGCAGGUGGGCAAAGAUGUCC3

B: 5’GGGTGAACCTGCAGGTGGGCAAAGATGTCCCAGCAAGGCACTGGTAGAATTCGGCAGCGT

L: DNA-5’GGACATCTTTGCCCACCTGCAGGTTCACCC3’

### R-loop unwinding assays

R-loop unwinding reactions (10 μL) contained 10 nM of R-loop, 1 mM ATP, 1-4 μM respective protein in R-loop buffer (6.6 mM Tris pH7.5, 3% glycerol, 0.1 mM EDTA, 1 mM DTT, 0.5 mM MgCl2) and were incubated at 37°C for 10min. Reactions were stopped by adding 2 μl of stop buffer (10 mg/ml proteinase K (NEB) and 1 % SDS) and incubated for 10 minutes at 37°C. 2 μl 50% glycerol was added to samples which were run on 6% PAGE gels at 100 V for 1 hr in 1 x TBE.

### Quantification and Statistical Analysis

Statistical significance for comparison of means was generally assessed by Student’s t test. All data shown is from at least 3 independent experiments ± standard error of the mean (s.e.m) unless indicated. Statistically significant pair-wise comparisons are indicated on each Figure where * refers to p <0.05; **, p <0.01; ***, p <0.001; ****, p <0.0001.

## Supplementary Figure Legends

**Figure S1:**
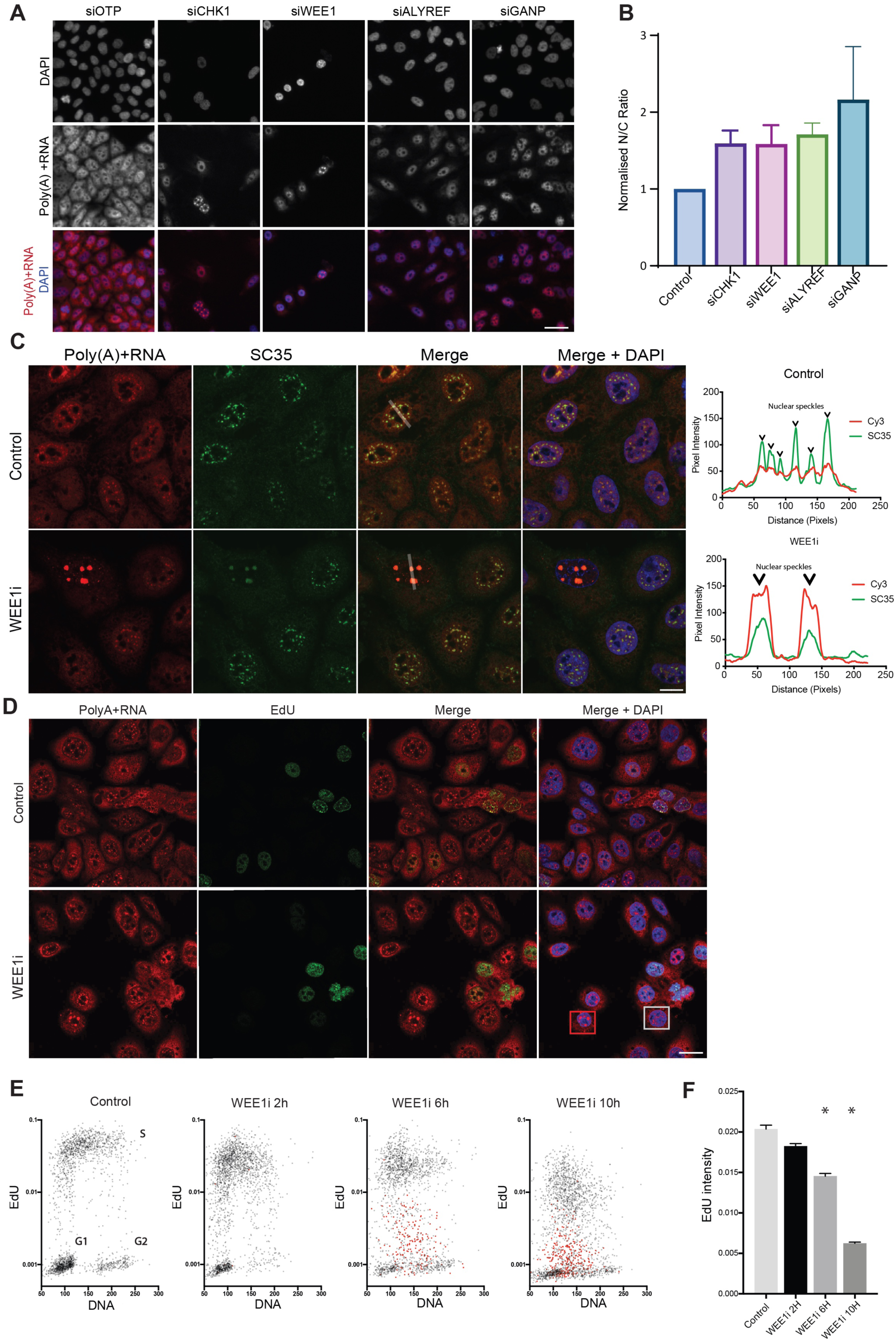
CRISPR and siRNA screens both identify cell cycle and DNA damage associated kinases WEE1 and CHK1 as mRNA export regulators. **(A**) Representative images from a high content imaging screen of poly(A)+RNA localisation. siRNA controls, positive controls ALYREF and GANP as well as novel regulators CHK1 and WEE1 are shown. (**B**) Normalised nuclear cytoplasmic ratio (N/C) ratio of poly(A)+RNA following GANP ALYREF, CHK1 and WEE1 depletion. The graph represents the mean from 3 experiments with error bars representing SD. (**C**) Representative images displayed from confocal microscopy of FISH for poly(A)+RNA and concomitant IF for SC35. Line scan fluorescence of the transparent white line in the merge channel is graphed to assess co-localisation. Nuclear speckles are highlighted by arrow heads. (**D**) Representative images displayed from confocal microscopy of FISH for poly(A)+RNA and Clickit-EdU-488. EdU was spiked in for the last 10 minutes prior to fixation. Cells with enlarged polyA(+) RNA foci are highlighted in red and a representative cell in highlighted in grey. (**E**) Representative plots of nuclear intensities of EdU and DNA (DAPI) in individual cells from n of 3 experiments for the indicated treatments. A minimum of 2000 cells were analysed. Cells with enlarged nuclear speckles are highlighted in red. (**F**) Mean EdU intensities measured from (**E**). Error bars represent SEM with * for p<0.05. The white line on all merge channels represents 25 µm.

**Figure S2:**
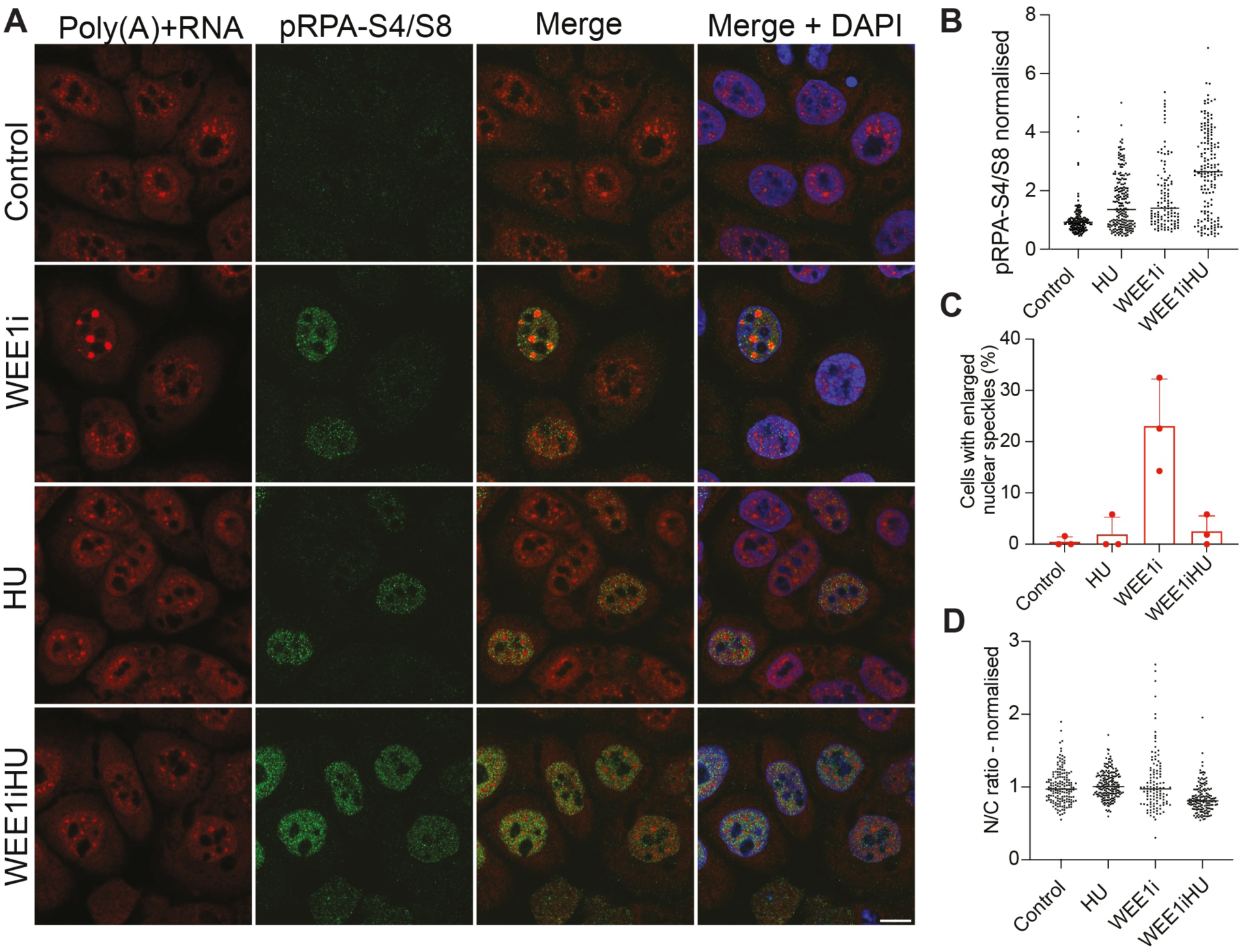
Cells with enlarged polyA(+)RNA nuclear speckles are rescued by treatment with replication stress inducer hydroxyurea. **(A**) Representative images displayed from confocal microscopy of FISH for poly(A)+RNA and pRPA-S4/S8. Cells with enlarged polyA(+)RNA foci are highlighted in red. **(B**) Nuclear pRPA-S4/S8 levels across three independent experiments with a minimum of 50 cells measured for the indicated treatments. Dots represent individual cells, and the line represents the mean. (**C**) Cells with nuclear speckles 1 standard deviation larger the mean of the control represented as a percentage for the indicated treatments. Individual dots represent the means of each experimental repeat (n=3) with error bars representing SD. **d**, N/C ratio of poly(A)+RNA for the indicated treatments. Individual dots represent the means of each repeat, and the line represents the mean.

**Figure S3:**
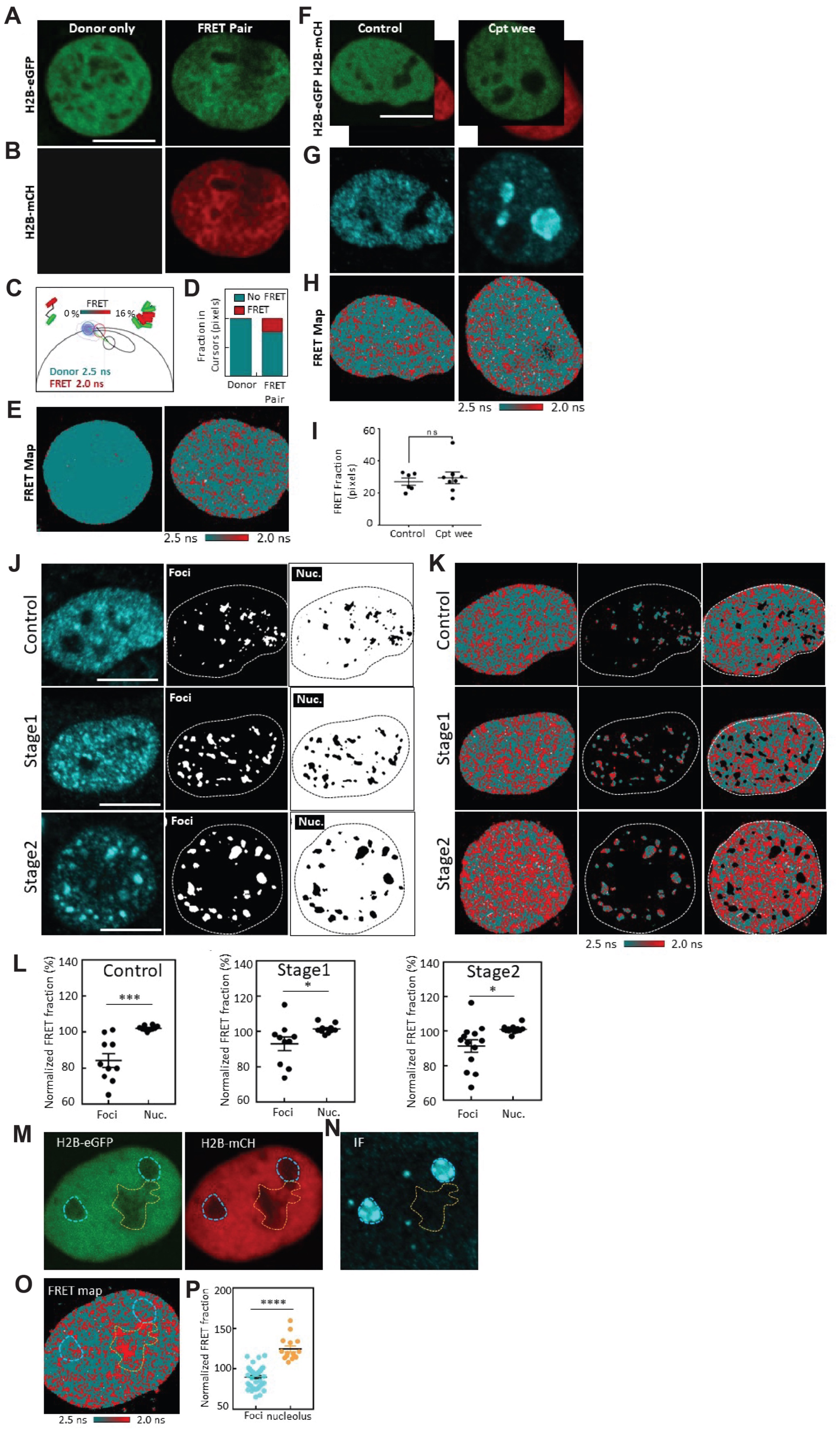
FLIM/FRET analysis of CAL51 cells following replication stress and WEE1 inhibition. (**A**) Fixed G_1_ Cal51 nucleus expressing H2B-eGFP in the absence of H2B-mCh (Scale bar 10 µm). (**B**) Fixed G_1_ Cal51 nucleus expressing H2B-eGFP in the presence (RIGHT) of H2B-mCh. (**C**) Phasor distribution of H2B-eGFP in the absence (donor control) versus presence of H2B-mCh (histone FRET experiment) with a theoretical FRET trajectory superimposed (black curve) that extends from the unquenched donor lifetime (teal cursor, 2.5ns). This FRET trajectory enables characterization of the histone FRET efficiency as 12% (red cursor, 2.1 ns) and definition of a palette to detect open (teal) versus compact (red) chromatin. (**D**) Quantification of the fraction of pixels in the phasor cursor that reports no FRET (open chromatin) versus histone FRET (compact chromatin) in the cells presented in panel (**A-B**). (**E**) FLIM maps of H2B-eGFP in the absence (LEFT) versus presence (RIGHT) of H2B-mCh pseudo-coloured according to the FRET palette defined in the phasor plot of panel (**C**). (**F**) Control (left) and camptothecin and WEE1 inhibitor treated G_1_ CAL51 cells co-expressing histone FRET pair (H2B-eGFP & H2B-mCh) (Scale bar 10 µm). (**G**) Immunofluorescence for SRRM2 in cells from (**F)**. (**H**) FLIM map of the cell presented in panel (**F-G**) pseudo-coloured to report histone FRET (red pixels) versus non-FRET (teal pixels). (**I**) Quantification of fraction of histone FRET in the across multiple cells in control and camptothecin and WEE1 inhibitor treated conditions (N= 6 or 7 cells, two biological replicates). The scatter plot shows individual cell value with SEM. ns *p* > 0.05 (unpaired *t*-test). (**J-L**) IF-based mask analysis of histone FRET reveals chromatin to be ‘open’ at sites of poly(A)+RNA nuclear speckles. (**J**) Representative SRRM2 IF image (left) in untreated, early (stage 1) and middle stage (stage 2) of camptothecin and WEE1 inhibitor treated G1 CAL51 cell nucleus that is co-expressing histone FRET pair (H2B-eGFP & H2B-mCh), masks based on SRRM2 IF intensity that select chromatin inside (middle) and outside (right) of poly(A)+RNA nuclear speckles. (**K**) Histone FLIM maps of the whole cell nucleus (left), inside (middle) and outside (right) of poly(A)+RNA nuclear speckles with threshold defined by masks presented in panel **J**. (**L**) Quantification the fraction of histone FRET inside and outside of normal poly(A)+RNA nuclear speckles, early and middle stage enlarged poly(A)+RNA nuclear speckles across multiple cells (N=10-13, two biological replicates). The scatter plot shows individual cell value with SEM. * *p* < 0.05, *** *p* < 0.001 (paired *t*-test). (**M**) Both enlarged poly(A)+RNA nuclear speckles (highlighted by blue dashed line) and nucleolus (highlighted by yellow dashed line) have low H2B-eGFP (left) and H2B-mCh (right) intensity. (**N**) SRRM2 IF image of the cell presented in panel (**M**). (**O**) Histone FLIM map of the cell presented in panel (**M**). (**P**) Quantification of FRET fraction inside of enlarged poly(A)+RNA nuclear speckles and nucleolus across multiple cells. (N=15, two biological replicates). The scatter plot shows individual cell value with SEM. \*\*\*\**p* < 0.0001 (unpaired *t*-test).

**Figure S4:**
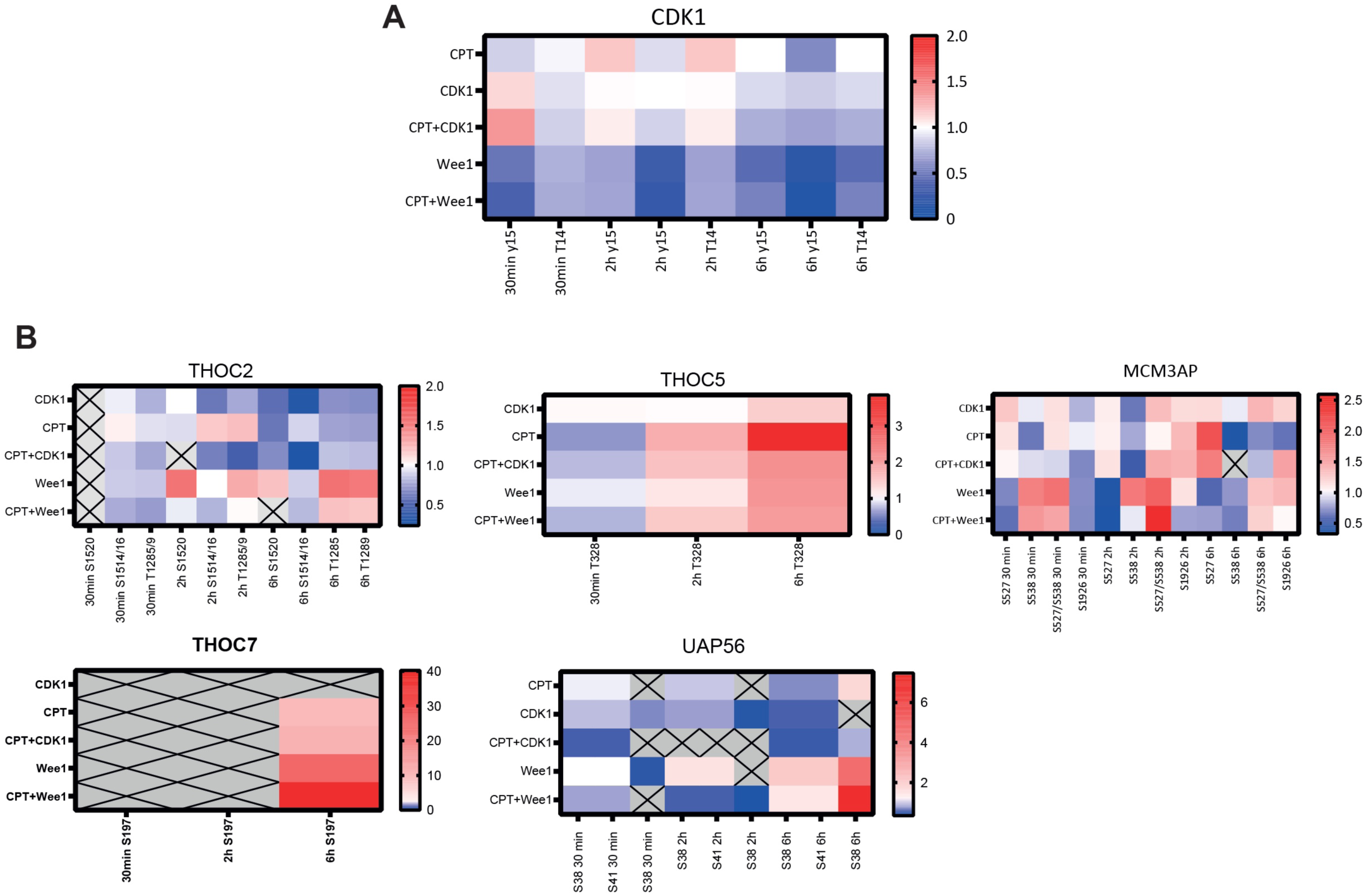
mRNA export factors are phosphorylated in response to replication stress. (**A**) Heat map of phosphorylation changes of CDK1 identified by the high throughput EasyPhos experiment presented in Figure 4. Colours represent a fold change of phosphorylation normalised to control. Peptides that were not detected (ND) in particular timepoints are represented in grey. (**B**) Heat map of phosphorylation changes of mRNA export factors THOC2, THOC5, MCM3AP, THOC7 and UAP56 identified by the EasyPhos experiment presented in Figure 4. Colours represent a fold change of phosphorylation-normalised to control.

**Figure S5:**
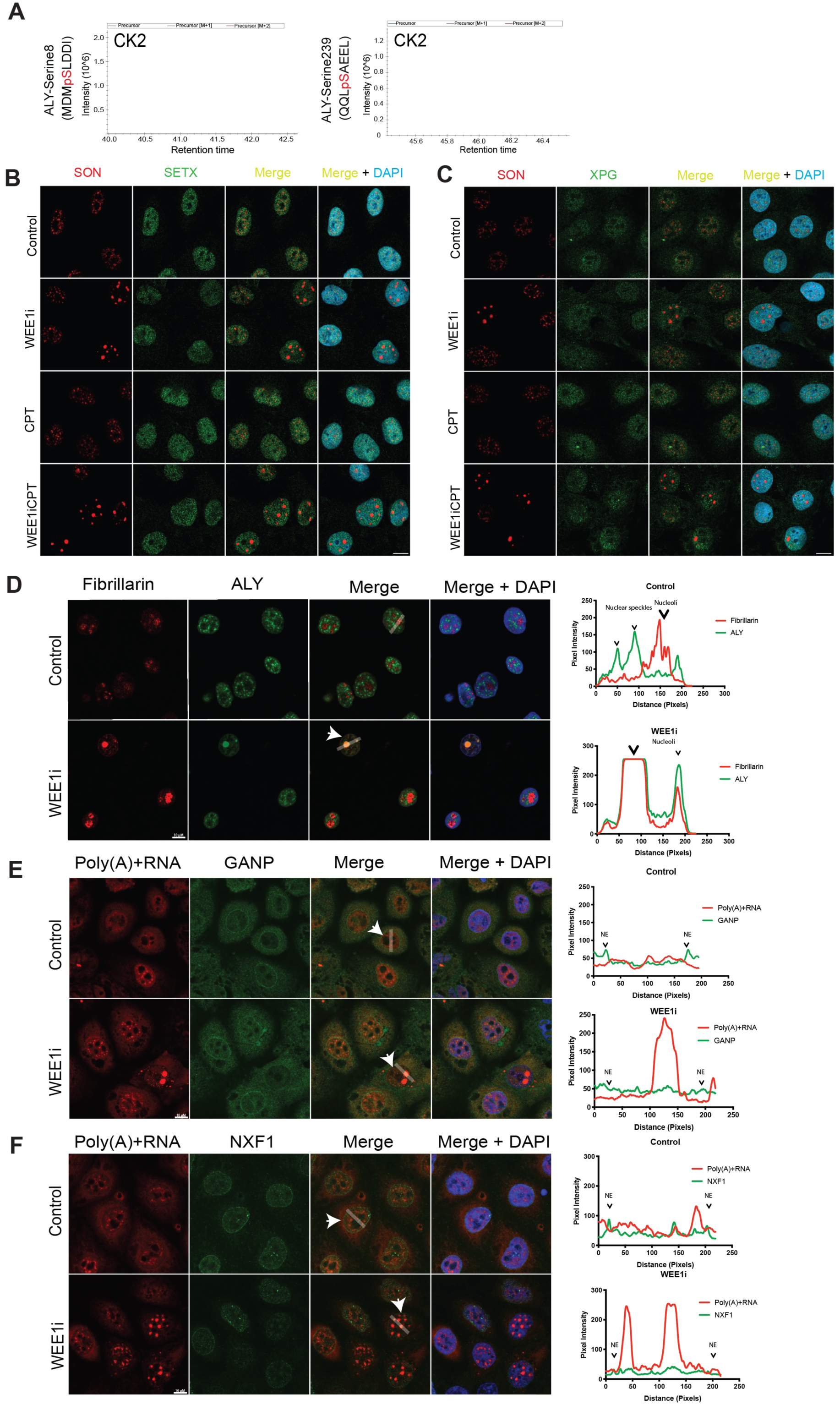
mRNA export factors are mis-localised from enlarged nuclear speckles in cells with high levels of replication stress to prevent nuclear export of mRNA. **(A)** *In vitro* kinase assays with CK2 and GST-ALYREF, from the same experiment in Figure 5. Peptides indicated were identified in Figure 5 as phosphorylated by CDK1 or PLK1. These peptides were not phosphorylated by CK2 as determined by mass-spectrometry. **(B**) Senataxin localisation is not altered following replication stress. Representative confocal microscopy images of dual IF for Senataxin and nuclear speckle marker SON for the indicated treatments are shown. (**C**) XPG localisation is not altered following replication stress. Representative confocal microscopy images of dual IF for XPG and nuclear speckle marker SON for the indicated treatments are shown. (**D**) Dual IF for ALYREF and nucleolar marker fibrillarin in cells treated with DMSO or WEE1 inhibition. Localisation was assessed by a line scan with areas interrogated displayed by the white arrow and line. (**E**) FISH for poly(A)+ RNA with concomitant IF for GANP in cells treated with DMSO or WEE1 inhibition. Localisation was assessed by a line scan with areas interrogated displayed by the white arrow and line. **(F**) FISH for poly(A)+ RNA with concomitant IF for NXF1 in cells treated with DMSO or WEE1 inhibition. Localisation was assessed by a line scan with areas interrogated displayed by the white arrow and line.

**Figure S6:**
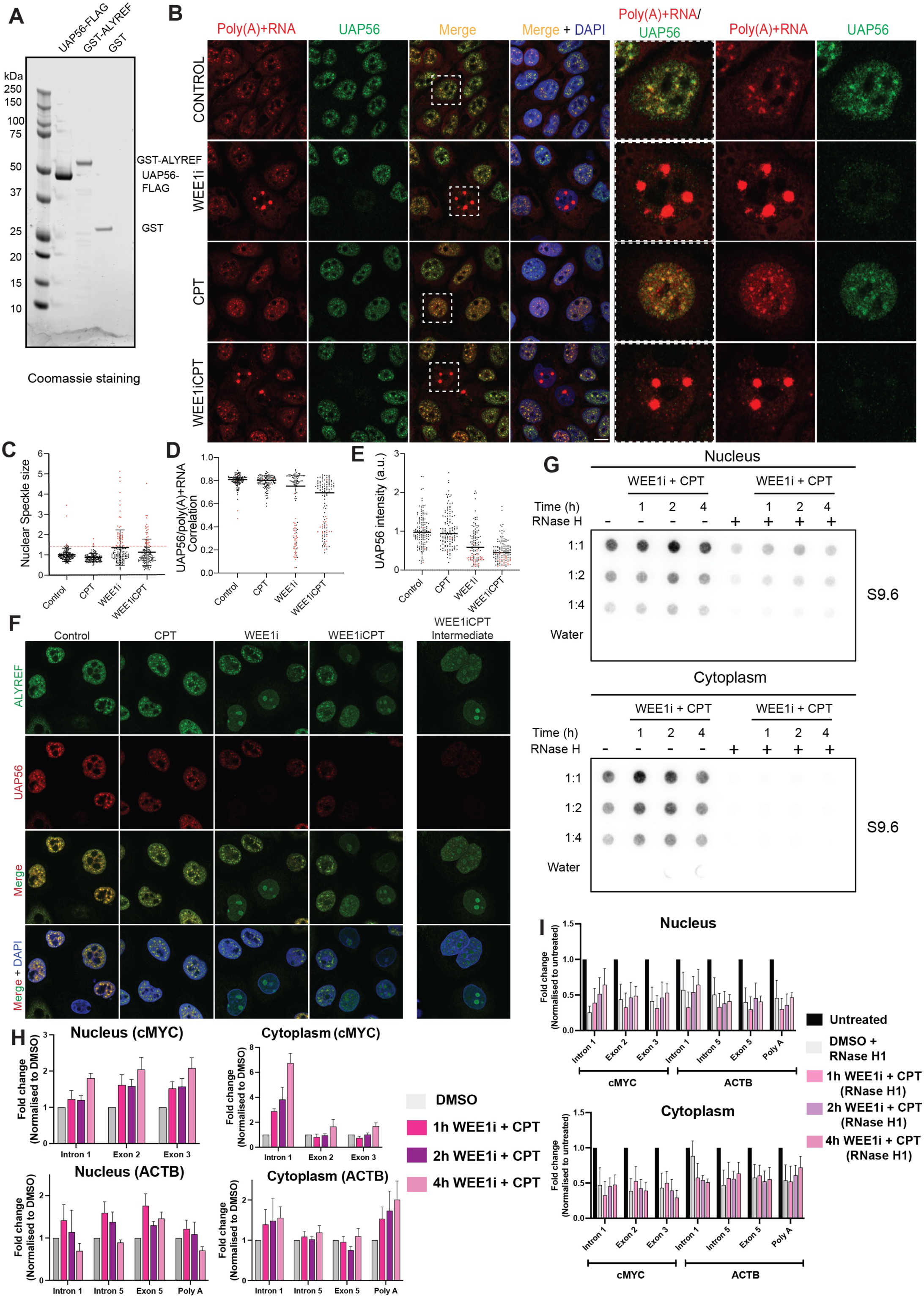
ALYREF and UAP56 co-operate to resolve R-loops, their co-localisation is altered following WEE1 inhibition and camptothecin treatment and R-loops can localise to the cytoplasm. (**A**) Coomassie staining for UAP56-FLAG, GST-ALYREF and GST proteins used for R-loop unwinding assays in Figure 6. (**B**) Representative images of FISH for poly(A)+ RNA with concomitant IF for UAP56 in double thymidine synchronised cells treated for 6 hours with the indicated inhibitor/s. Individual cells from the merge channel are selected and displayed at a higher magnification, those selected are indicated by the white box. (**C**) Nuclear speckle size analysed from (**B)** with a minimum of 50 cells analysed per experimental repeat (n=3). Cells with nuclear speckles greater than one SD above the control are highlighted in red. (**D**) UAP56/Poly(A)+RNA correlation from (**B)** with a minimum of 50 cells analysed per experimental repeat (n =3). Cells with nuclear speckles greater than one SD above the control are highlighted in red. (**E**) Nuclear UAP56 IF intensity analysed from B with a minimum of 50 cells analysed per experimental repeat (n =3). Cells with nuclear speckles greater than one SD above the control are highlighted in red. (**F**) Dual IF for ALYREF and UAP56 in double thymidine synchronised cells treated for 6 hours with the indicated inhibitor/s. An image displaying an intermediate phenotype of partial ALYREF/UAP56 mis-localisation for WEE1i and camptothecin treatment is also presented. (**G**) Nuclear and cytoplasmic R-loop levels are increased following WEE1 inhibitor and Camptothecin treatment. A representative dot blot using the S9.6 antibody to detect R-loops is shown following treatment with a WEE1 inhibitor and Camptothecin treatment for the indicated times. Some samples were treated with RNase H prior to the pulldown experiment. (**H**) DRIP qPCR assessment of R-loop formation in cMYC and ACTB genes in the indicated genomic regions from nuclear and cytoplasmic fractions following WEE1 inhibitor and camptothecin treatment. Graphs represent the mean of three independent experiments with error bars showing SEM. (**I**) Samples from (**H**) were also treated with RNase H. Graphs represent the mean of three independent experiments with error bars showing SEM.

**Figure S7:**
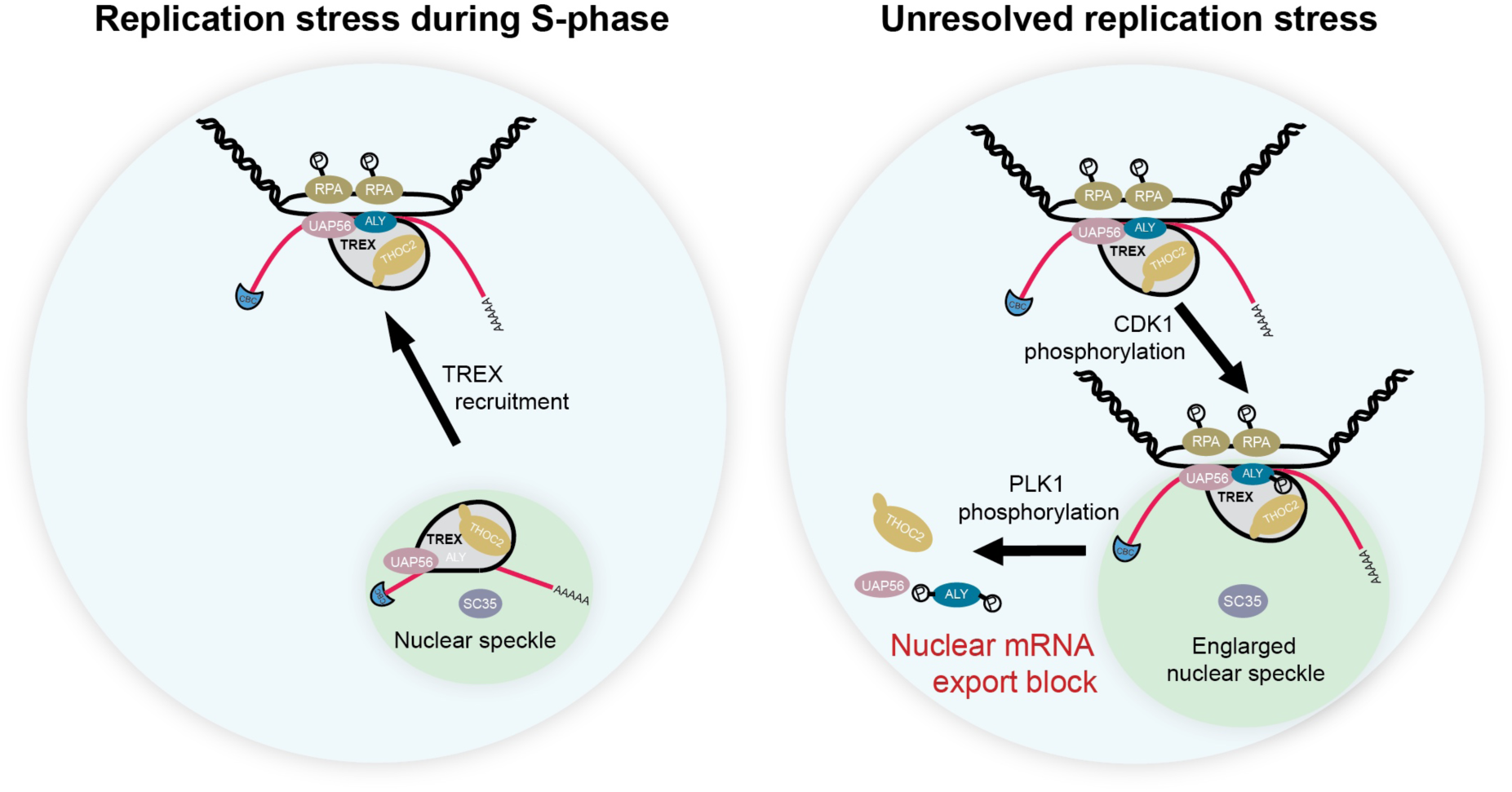
Model for how mRNA export factors are regulated during replication stress in S-phase. mRNA export factors are recruited to sites of replication stress for resolution of R-loops by ALYREF and UAP56 (left panel). If replication stress is unresolved (right panel), ALYREF phosphorylation at serine 239 by CDK1 can drive re-localisation of poly(A)+RNA and R-loops to nuclear speckles for resolution. If R-loops cannot be resolved by UAP56 and ALYREF at nuclear speckles, ALYREF can then be phosphorylated by PLK1 at serine 8, which can abolish the interaction between UAP56 and ALYREF promoting their re-localisation from nuclear speckles and preventing export of R-loop associated mRNA from occurring.

